# Compressed Cortical Input Separates Control from Dynamics in Striatum

**DOI:** 10.1101/2025.09.05.674585

**Authors:** Sreejan Kumar, Matthieu B. Le Cauchois, Alexander Mathis, Lea Duncker, Jonathon R. Howlett, Marcelo G. Mattar

## Abstract

The dorsolateral striatum (DLS) supports diverse time-sensitive behaviors—action chunking, duration estimation, and motor timing—yet no single framework is able to explain all of these phenomena. Here, we propose that the massive convergence of cortical projections onto the striatum provides such a framework. We develop a corticostriatal neural network model in which a recurrent cortical module communicates with a recurrent striatal module through a low-dimensional, noisy bottleneck, with the whole system trained via reinforcement learning. Across three DLS-associated tasks, compression produces a consistent computational motif whereby cortex provides low-dimensional control signals while the striatum generates stable, time-encoding dynamics. This separation gives rise to chunking behavior with action slipping, intensity-biased duration judgements with stimulus-modulated time coding, and stereotyped motor timing programs. Perturbation of the compressed cortical signal causally shifts behavior while preserving sequential structure in striatum activity. In sum, our results unify information-theoretic and dynamical systems perspectives of basal ganglia function to link anatomical compression to a variety of temporally-sensitive sensorimotor behaviors implicated in the DLS.

## Introduction

The dorsolateral striatum (DLS), the sensorimotor input territory of the basal ganglia, has been implicated in a remarkably broad range of time-sensitive behaviors: the acquisition and execution of action sequences (Graybiel, 1998; X. Jin & Costa, 2015; Martiros et al., 2018), the estimation and comparison of elapsed durations (Gouvêa et al., 2015; Monteiro et al., 2023; Rodrigues et al., 2024; Toso et al., 2021), the temporal coordination of learned motor habits (Dhawale et al., 2021; Hidalgo-Balbuena et al., 2019; Rueda-Orozco & Robbe, 2015), and more broadly, the selection of actions appropriate to the current moment within a temporally extended behavioral context (D. Z. Jin et al., 2009; Klaus et al., 2017; Markowitz et al., 2023). A unifying theme across these functions is the need to produce the right action at the right time (Haimerl et al., 2025). Yet despite decades of work exploring each of these phenomena individually, a computational framework explaining why a single brain region supports such diverse temporally-sensitive functions has remained elusive. Here, we propose that the answer lies in a defining anatomical feature of the corticostriatal pathway: the massive convergence of cortical projections onto the striatum.

Neural projections from the cortex to the striatum undergo a dramatic dimensionality reduction: the number of cortical neurons projecting to the striatum exceeds the number of receiving striatal neurons by more than an order of magnitude (Figure 1A). The principal neurons of the striatum, medium spiny neurons (MSNs), which comprise approximately 95% of the striatal population (Kita & Kitai, 1988), possess large, extensively branching dendritic arbors studded with spines that receive convergent excitatory input from thousands of cortical neurons (Kincaid et al., 1998; Wilson & Groves, 1980). Bar-Gad, Morris, & Bergman (2003) termed the resulting population-level compression “anatomical funneling” and showed that information transmitted through the basal ganglia is decorrelated relative to cortical input, consistent with dimensionality reduction at the corticostriatal interface. Their theoretical framework of “reinforcement-driven dimensionality reduction” (RDDR) proposed that this compression is optimized for reward-relevant dimensions — akin to PCA, but with a loss function defined by reinforcement rather than variance explained (Bar-Gad, Havazelet-Heimer, et al., 2003). However, RDDR characterizes corticostriatal compression in static, information-theoretic terms; it does not address how this compression shapes the *dynamics* of the downstream circuit.

**Fig. 1.**
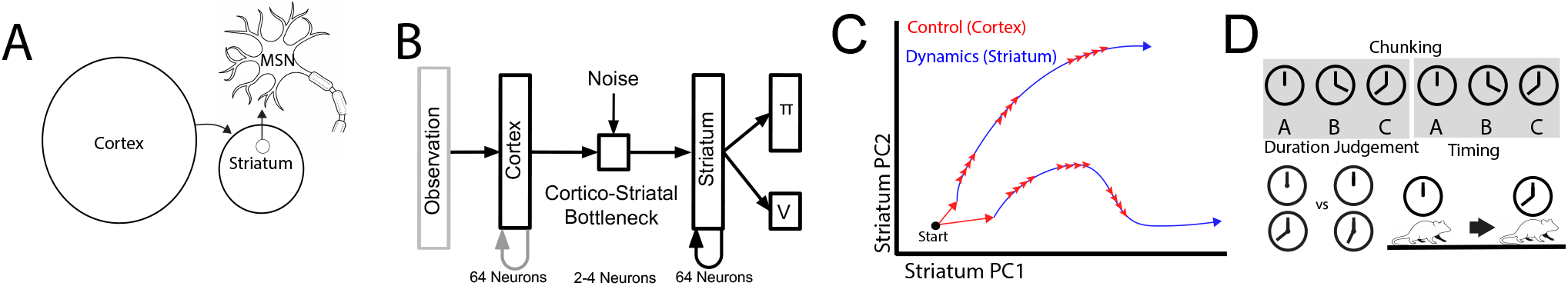
Corticostriatal Compression Framework. (**A**). **Anatomical motivation**. Cortical neurons projecting to the striatum outnumber receiving striatal neurons by more than an order of magnitude. Medium spiny neurons (MSNs), the principal striatal cell type, possess large branching dendrites that integrate convergent input from diverse cortical sources. (**B**). **Model architecture**. A cortical RNN (64 units) communicates with a striatal LSTM (64 units) through a low-dimensional noisy bottleneck (2–4 units). The striatal module produces a policy (*π*) and value estimate (*V*), trained via reinforcement learning. Cortico-cortical weights are random and fixed; all striatal and cortico-striatal weights are plastic. (**C**). **Conceptual schematic**. Compression forces a separation of low-dimensional control signals (cortex, red) from stable time-encoding dynamics (striatum, blue). Cortical input guides progression along trajectories generated by the striatum. (**D**). **Task domains** Three tasks used to test the framework: sensorimotor chunking, duration judgements, and habitual motor timing, all associated with the DLS.

More recent work has shifted focus from the static properties of corticostriatal compression to the dynamics of the striatal circuit itself. Striatal population activity evolves along reproducible trajectories in neural state space, and the timecourse of this evolution provides a temporal basis for the computations the striatum performs (Monteiro et al., 2023; Tsao et al., 2022). Cortical and thalamic input appears to parametrically control these dynamics — modulating the speed and gain of striatal trajectories — while the striatum retains autonomy over the structure of the activity patterns themselves (Hidalgo-Balbuena et al., 2019; Lee et al., 2019; Murray & Escola, 2017). Yet this dynamical systems perspective has developed largely independently of the information-theoretic account of corticostriatal compression, and the two have rarely been connected within a single computational framework. A framework that bridges these ideas could explain how a single anatomical motif simultaneously gives rise to both the behavioral and dynamical signatures observed across DLS-dependent tasks.

We propose that corticostriatal compression is precisely this bridging principle: dimensionality reduction at the corticostriatal interface forces a separation of function in which the cortex provides low-dimensional control signals while the striatum generates intrinsic stable time-encoding dynamics, and it is this separation that gives rise to the structured temporal behaviors associated with the DLS (Figure 1C).

To instantiate this hypothesis, we developed a multiregion recurrent neural network model inspired by the architecture of Mizes et al. (2024), who modeled the corticostriatal pathway as a cortical module projecting to a recurrent striatal module, with sensory cues provided to cortex but not directly to the striatum (reflecting their previous finding that DLS activity can be insensitive to specific sensory cues, Mizes et al. 2023). Our model (Figure 1B) consists of a cortical module (a recurrent neural network with 64 units) that communicates with a striatal module (an LSTM with 64 units) through a low-dimensional noisy bottleneck (2–4 units). The striatal module produces both a policy and a value estimate, and the system is trained end-to-end via reinforcement learning using an actor-critic framework. Cortical recurrent weights are randomly initialized and frozen (Buonomano & Maass, 2009; Dominey, 1995), reflecting the assumption that cortical dynamics are shaped by a separate learning signal operating on a different timescale (Doya, 2000; Pasupathy & Miller, 2005), while all striatal and corticostriatal weights are plastic. The striatal module is recurrent, reflecting inhibitory interactions among MSNs and excitatory re-entrant loops through the basal ganglia output nuclei and thalamus (Lanciego et al., 2012; Mandelbaum et al., 2019; McElvain et al., 2021), as also modeled by Mizes et al. (2024). Whereas Mizes et al. (2024) imposed a penalty on the strength of the cortical-to-striatal projection, implicitly constraining information flow, our model makes this constraint explicit by directly limiting the dimensionality of the corticostriatal channel — enabling us to systematically study the functional consequences of compression. Our architecture thus captures the spirit of RDDR — cortical information is compressed through a capacity-limited channel optimized by reinforcement — while simultaneously implementing a dynamical system in which compressed cortical input parametrically controls the temporal evolution of striatal trajectories.

We test this model on three time-sensitive sensorimotor behaviors associated with the DLS (Figure 1D): a sensorimotor chunking task inspired by human behavioral experiments (Lai et al., 2025), a duration comparison task based on recordings from rat DLS (Rodrigues et al., 2024; Toso et al., 2021), and a motor timing task (Hidalgo-Balbuena et al., 2019). Across all three tasks, compression produces a consistent computational motif: the cortex provides low-dimensional, task-relevant control signals while the striatum intrinsically generates stable, time-encoding dynamics. This separation gives rise to human-like chunking with action slipping, intensity-biased duration judgements with stimulus-modulated time coding, and stereotyped motor timing programs with perturbation-sensitive timing. In each domain, perturbation of the compressed cortical signal causally shifts behavior while preserving the sequential structure of striatal activity. Our results thus provide a unifying mechanistic link between corticostriatal compression and the diverse temporally-sensitive functions of the DLS, bridging the information-theoretic and dynamical systems frameworks that have each been used separately to study this circuit.

## Results

### Sensorimotor Chunking

We first asked whether corticostriatal compression can account for sensorimotor chunking, a core DLS-associated behavior in which frequently co-occurring actions are grouped into single behavioral routines. To test this, we trained our model on a task inspired by Lai et al. (2025) in which participants observe sequences of stimuli drawn from a set of 16 possible items and must select the corresponding action for each stimulus to receive a reward (Figure 2A,B). The key manipulation is that eight of these stimuli (S0–S7, “chunk stimuli”) frequently appear in a fixed temporal order, forming a “chunk” during structured episodes, while the remaining “non-chunk” stimuli always appear in random order. Although chunk stimuli can appear individually in random contexts, they more frequently co-occur together in their fixed sequence via structured episodes (see Methods for full task parameterization). This design creates an opportunity for behavioral policy compression: rather than treating each stimulus independently, an efficient learner can recognize and exploit the predictable chunk sequence.

**Fig. 2.**
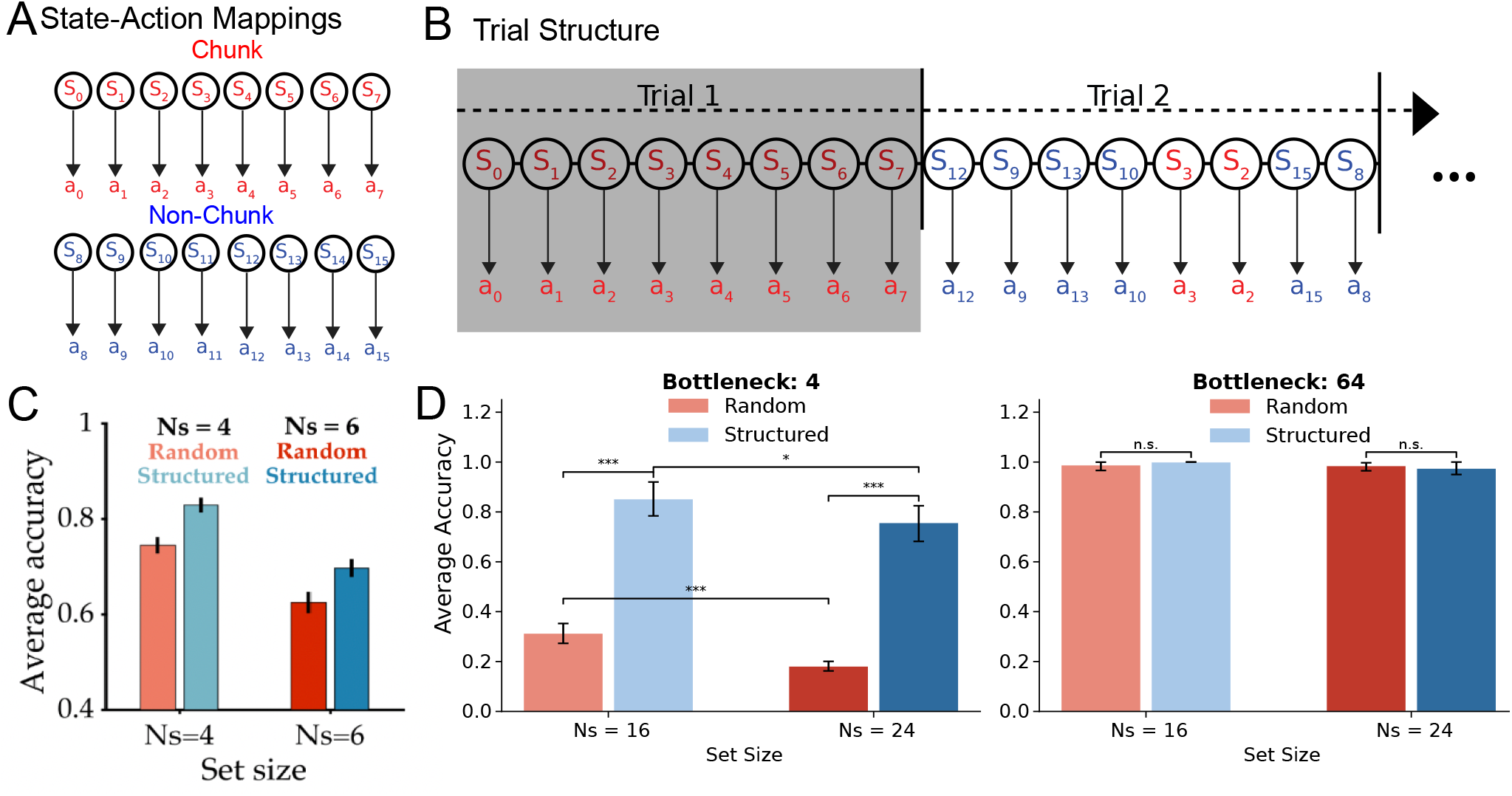
Corticostriatal Compression Reproduces Human-like Chunking. (**A**). **Sensorimotor chunking task**. Each stimulus has one rewarded action; the number of possible actions equals the number of stimuli. (**B**). **Task structure**. A sequence of *T* stimuli is sampled via either a structured or random scheme. In structured episodes (shaded), a fixed subset of stimuli appears in a predictable order (the “chunk”). In random episodes (unshaded), stimuli are drawn randomly, with chunk stimuli appearing at lower probability. (**C**). **Human behavioral signatures of chunking**. Data from Lai et al. (2025). Performance decreases with larger set sizes (*N*_*s*_ = 4 vs. *N*_*s*_ = 6) and improves on structured versus random trials (red vs. blue). (**D**). **Only strong compression reproduces both human signatures**. The 4-dimensional bottleneck (left) shows a significant structured-trial advantage at both set sizes (one-sided Welch’s *t*-test, both *p <* 0.001) and a set-size effect for both structured (*p* = 0.044) and random (*p <* 0.001) trials. The 64-dimensional bottleneck (right) shows neither effect (both *p >* 0.15). Error bars: 95% CI across 11 training seeds.

Human performance on this task reveals two signatures of capacity-limited processing that serve as qualitative targets for our model. First, performance decreases as the total number of stimuli increases, reflecting the cognitive cost of maintaining more stimulus-response mappings (Figure 2C, light vs. dark bars). Second, humans perform significantly better on structured trials containing the chunk than on random trials, demonstrating that they successfully exploit the temporal regularity (Figure 2C, red vs. blue bars). These two effects — degraded performance with increased memory load and improved performance with predictable structure — are hallmarks of policy compression in capacity-limited systems (Lai & Gershman, 2021).

To test whether corticostriatal compression could reproduce these behavioral signatures, we trained models with varying bottleneck dimensions and evaluated performance on both structured and random trials across set sizes, matching the conditions used in the human experiments. The uncompressed model (64-dimensional bottleneck) performed equally well across all conditions, showing neither the set size effect nor the structured trial advantage (Figure 2D, right), indicating that without a communication bottleneck, the model simply memorizes all stimulus-response mappings independently. By contrast, the compressed model (4-dimensional bottleneck) reproduced both human signatures: performance decreased with set size and improved on structured versus random trials (Figure 2D, left). These results demonstrate that corticostriatal compression is sufficient to produce human-like chunking behavior.

#### Structured Behavior and Striatal Time-Encoding Dynamics

If the compressed model genuinely chunks actions rather than simply showing a statistical advantage on structured trials, it should exhibit “action slipping” — the classic chunking phenomenon in which a familiar sequence is executed inflexibly even when inappropriate (Norman, 1981). We tested this by presenting partial chunk initiations (e.g., stimuli S0–S2) followed by random stimuli requiring different actions. The compressed model inflexibly completed the entire chunk sequence, ignoring subsequent non-chunk stimuli (Figure 3A, top), whereas the uncompressed model remained responsive to the actual stimuli presented (Figure 3A, bottom). This action slipping confirms that the compressed model has consolidated the chunk into a single behavioral unit that, once triggered, runs to completion.

**Fig. 3.**
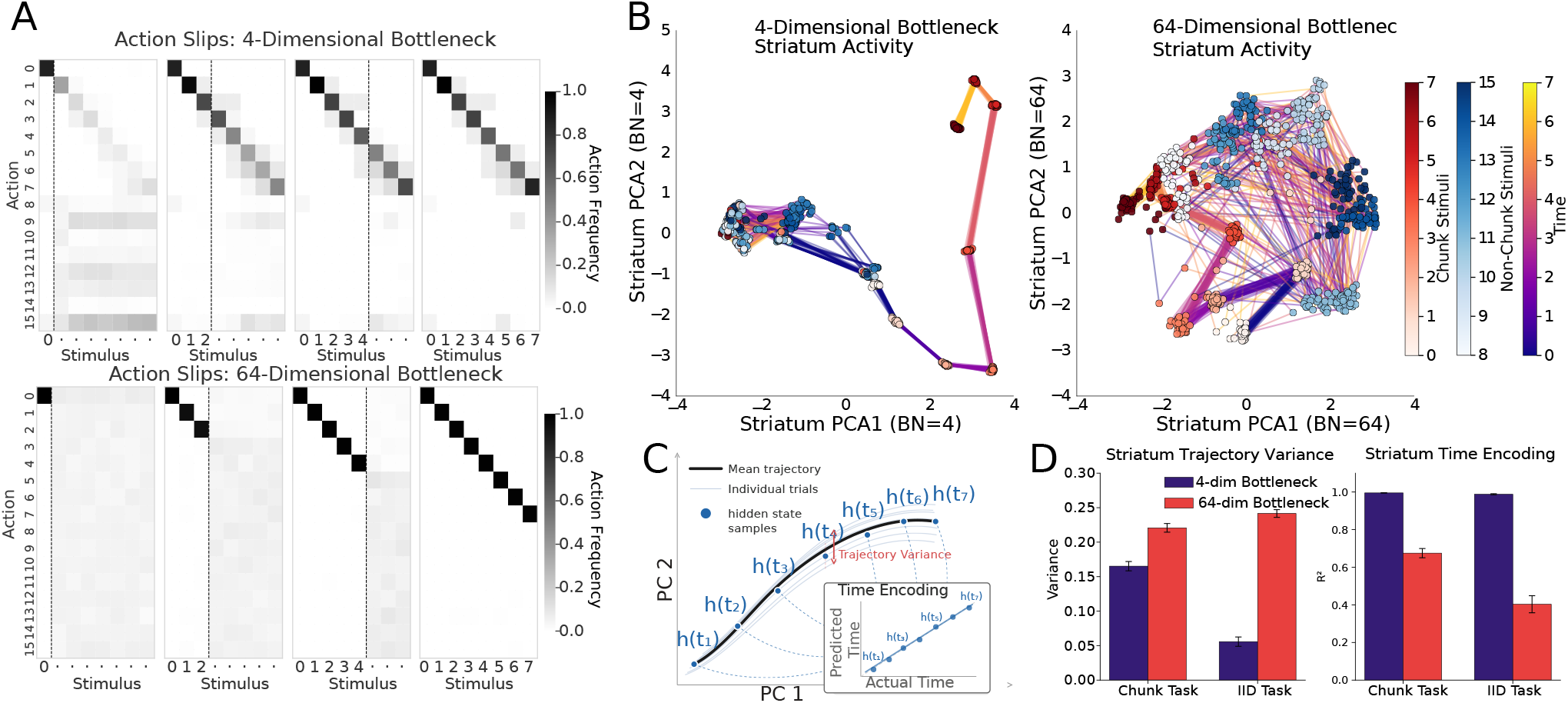
Corticostriatal Compression Produces Action Slipping and Stable Time-Encoding Striatal Dynamics. (**A**). **Action slipping**. Action distributions following partial chunk initiations (·: marginalization over future states; dashed lines: end of partial initiation). The compressed model (top) inflexibly completes the chunk sequence; the uncompressed model (bottom) remains responsive to actual stimuli. (**B**). **Striatal hidden state trajectories (PCA)**. Points colored by stimulus identity (chunk vs. non-chunk); lines colored by timestep. The compressed model (left, bottleneck dim = 4) develops a smooth, stereotyped trajectory for the chunk; the uncompressed model (right, dim = 64) shows stimulus-specific clusters. (**C**). **Quantification schematic**. Trajectory variance: trial-to-trial variability of the full hidden state at each timestep. Time encoding: cross-validated *R*^2^ of a linear decoder predicting elapsed time from hidden state. (**D**). **Trajectory variance and time encoding**. Compression reduces trajectory variance (left) and increases time encoding (right) in both the original task and an IID control (all *p <* 0.001, two-sided *t*-tests, *n* = 11 seeds). Error bars: 95% CI.

Corticostriatal compression reduces the sensory information available to the striatum, which should force the network to rely more on internally generated temporal dynamics. Consistent with this reasoning, stimulus identity was significantly harder to decode from the hidden state in models with smaller bottlenecks (Figure S1), suggesting that the striatum must rely more on internally generated activity patterns. We therefore examined the structure of the striatum’s dynamics directly using PCA of the hidden state trajectories (Figure 3B). Importantly, both the compressed and uncompressed models have the same striatal dimensionality (64 units); they differ only in the dimensionality of the input the striatum receives from cortex. In the compressed model, chunk stimuli evolved along a consistent, smooth trajectory while non-chunk stimuli collapsed into a single cluster, with position along the trajectory corresponding to elapsed time. By contrast, the uncompressed model’s striatum showed fragmented, stimulus-specific clusters with less temporally structured trajectories.

To quantify these observations, we developed two metrics computed on the full 64-dimensional hidden state (Figure 3C): *trajectory variance*, the mean trial-to-trial variability of the hidden state at each timestep, and *time encoding*, the cross-validated *R*^2^ of a linear decoder trained to predict elapsed time from the hidden state. Across both the chunking task and a control task where stimuli were drawn fully IID, models with corticostriatal compression showed significantly lower trajectory variance and significantly higher time encoding than models without compression (Figure 3D; all comparisons *p <* 0.001, two-sided *t*-tests, 11 seeds per condition). The IID result is particularly noteworthy: this environment contained no external temporal structure whatsoever, yet the compressed model still produced stable, time-encoding trajectories. These findings held when trial lengths were varied randomly, dissociating absolute time from task progress (Figure S2), and were robust to using a vanilla RNN instead of an LSTM for the striatal module (Figure S3). Together, these results confirm that corticostriatal compression produces stable time-encoding striatal dynamics across a range of task and architectural conditions.

A particularly striking result emerged from models that had never been trained. Random-weight models with a corticostriatal bottleneck already exhibited strong time-encoding trajectories, whereas random-weight models without a bottleneck did not (Figure S4). Both models use the exact same recurrent architecture and differ only in the dimensionality of the striatum’s input: the bottleneck model’s cortical input is randomly projected into a lower-dimensional space, whereas the non-bottleneck model retains the full cortical dimensionality. When input dimensionality is sufficiently constrained, the variance of the hidden state is dominated by the network’s internal dynamics rather than by input-driven variability, producing stable trajectories that implicitly encode time. This finding demonstrates that the capacity for temporal encoding arises from corticostriatal compression itself, existing independently of and prior to task-specific learning.

#### Control of Behavior from Compressed Cortical Input

Having established that corticostriatal compression produces both chunking behavior and stable time-encoding striatal dynamics, we next asked whether the compressed cortical representation causally controls chunk execution. If the bottleneck carries a low-dimensional control signal that triggers chunking, then artificially shifting the bottleneck along a chunk-associated dimension should induce chunk-like behavior even in the absence of structured input.

We first confirmed that chunk and non-chunk stimuli occupy separable regions of bottleneck space using PCA (Figure 4A). We defined the “chunk direction” as the vector from the non-chunk centroid to the chunk centroid, and applied additive perturbations of varying magnitude along this direction during random episodes — trials in which stimuli are randomly drawn and thus never contain the chunk sequence (Figure 4B). In the compressed model (4-dimensional bottleneck), perturbation along the chunk direction induced a graded increase in correctly timed chunk actions per episode (Figure 4C). Critically, perturbations along random orthogonal directions in the same bottleneck space had no such effect, confirming direction specificity. The uncompressed model (64-dimensional bottleneck) showed a weaker effect along the chunk direction, consistent with the control signal being less consolidated in the absence of compression. These results demonstrate that corticostriatal compression consolidates chunk identity into a single direction in the bottleneck — a low-dimensional cortical control signal sufficient to drive the striatum’s intrinsic dynamics to produce a structured, temporally extended action sequence.

**Fig. 4.**
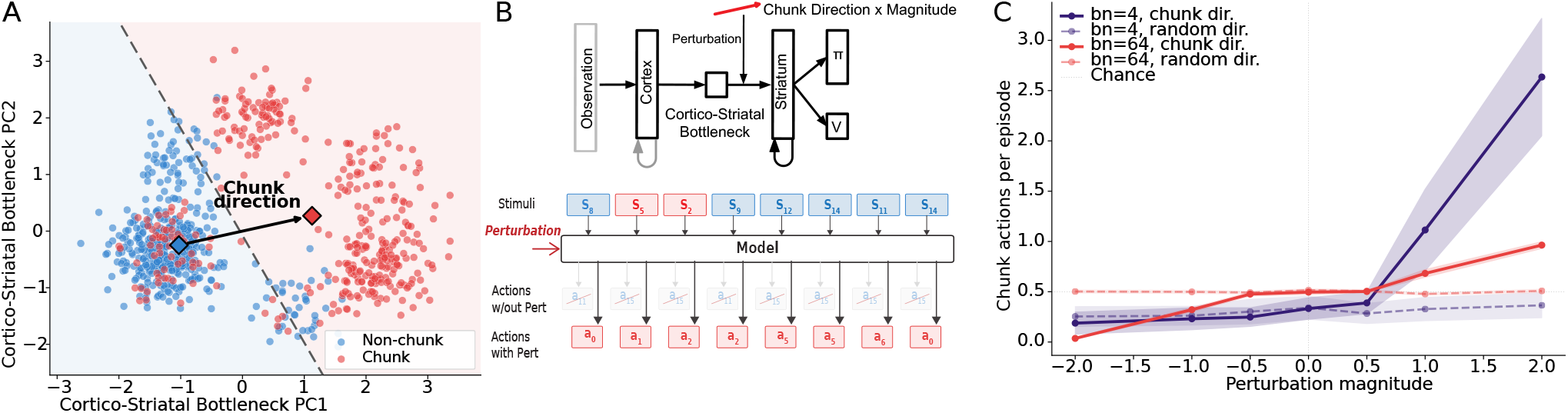
Compressed Cortical Activity Controls Chunking Behavior. (**A**). **Bottleneck PCA**. Chunk (red) and non-chunk (blue) stimuli are linearly separable in the 4-dimensional bottleneck space. (**B**). **Perturbation schematic**. During random trials, the bottleneck receives additive input along the chunk direction. Example trial: the perturbation induced 5/8 correctly timed chunk actions despite random stimulus presentation. (**C**). **Perturbation results**. Correctly timed chunk actions per episode increase with perturbation magnitude along the chunk direction for the compressed model (slope = 0.557, *p* = 4.4 *×* 10^−3^) but not along random orthogonal directions. The uncompressed model shows a weaker effect (slope = 0.212); random directions remain flat. SEM bands across *n* = 11 seeds.

### Duration Judgments

If the separation of cortical control from striatal dynamics is a general principle of corticostriatal function, it should extend beyond chunking to other time-sensitive behaviors associated with the DLS. We next tested our framework on a duration discrimination task in which Rodrigues et al. (2024), re-analyzing data from Toso et al. (2021), showed that DLS time-encoding dynamics are modulated by stimulus intensity and that this modulation systematically biases temporal judgements.

#### Structured Behavior and Striatal Time-Encoding Dynamics

In this task, Toso et al. (2021) trained rats to compare two sequential vibrotactile stimuli (*S*_1_, *S*_2_), each varying independently in both duration (*T*_1_, *T*_2_) and intensity (*I*_1_, *I*_2_), and to report which stimulus was longer (Figure 5A). Although intensity was irrelevant to the task, the analyses by Rodrigues et al. (2024) revealed that rats’ duration judgements were systematically biased by the intensity of the second stimulus: higher *I*_2_ led rats to report *T*_2_ as longer, as if intense stimuli were perceived as lasting longer. Critically, this behavioral bias had a neural correlate in the DLS — decoded time estimates during *S*_2_ were modulated by stimulus intensity, with higher intensity producing faster progression of the neural time estimate and thus inflating perceived duration. We simulated this task by training our model on duration comparison with Poisson spiking observations, where stimulus intensity modulated firing rate (see Methods). This entanglement is critical: because total spike count scales with both intensity and duration, the two features become ambiguous, creating the conditions under which intensity can bias temporal judgements.

**Fig. 5.**
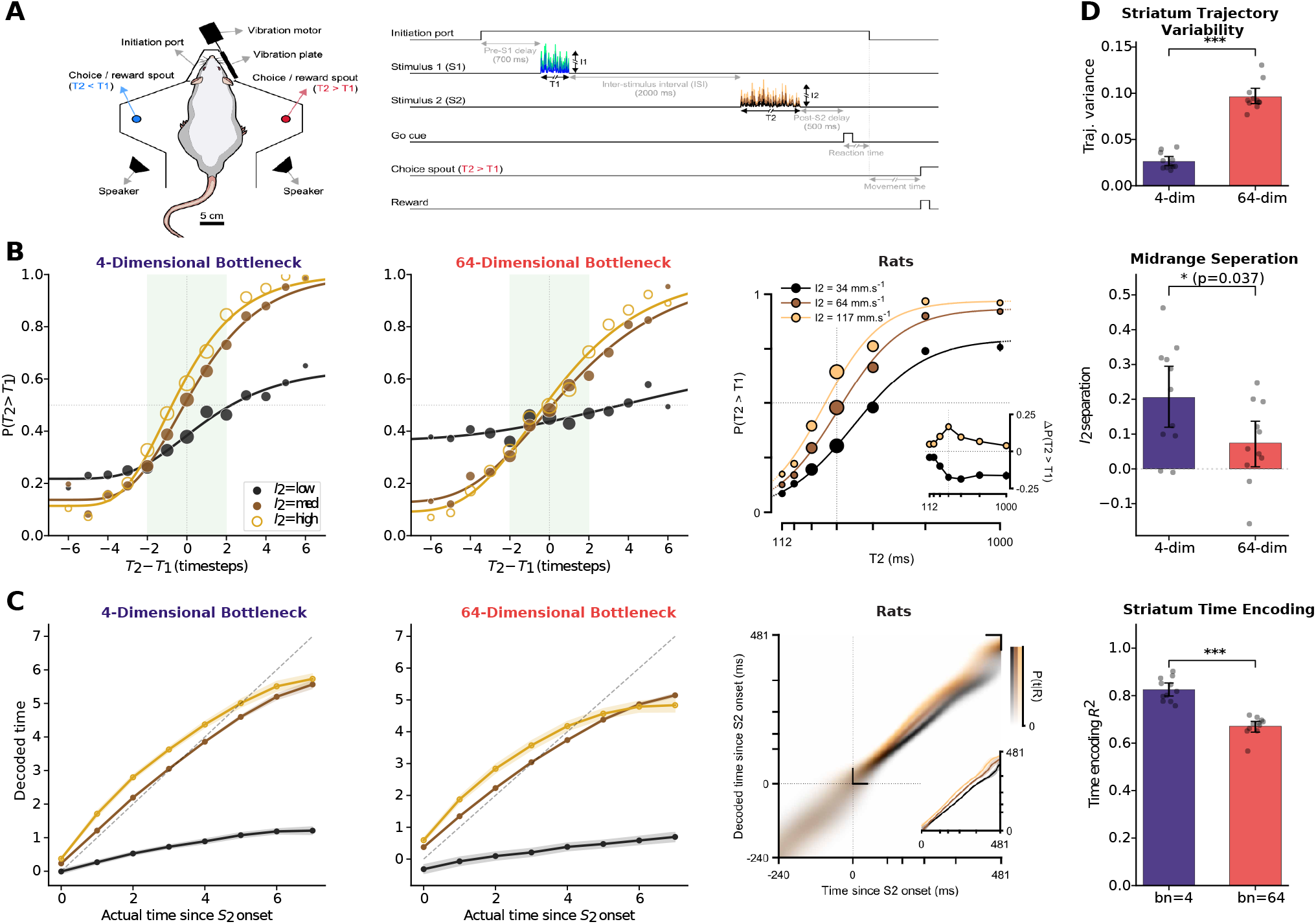
Corticostriatal Compression Reproduces Intensity-Biased Duration Judgements and Stable Striatal Time Encoding. (**A**). **Duration comparison task**. Rats compare two sequential vibrotactile stimuli on duration (*T*_1_, *T*_2_), with intensity (*I*_1_, *I*_2_) being task-irrelevant (Rodrigues et al., 2024; Toso et al., 2021). (**B**). **Psychometric curves split by** *I*_2_. The compressed model (left) shows clear intensity-dependent separation: high *I*_2_ biases reports toward “*T*_2_ longer.” The uncompressed model (middle) shows weaker separation. Right: rat behavioral data from Rodrigues et al. (2024). Far right: midrange separation (psychometric curve distance where |*T*_2_−*T*_1_| *≤*2) is significantly stronger in the compressed model (0.206 *±* 0.148 vs. 0.074 *±* 0.111; *p* = 0.037, *n* = 11 seeds). (**C**). **Striatal time decoding during** *S*_2_, **split by** *I*_2_. Decoded time progresses faster for high-intensity stimuli in both models. Right: analogous neural data from Rodrigues et al. (2024). Far right: overall time encoding *R*^2^ is stronger with compression (0.826 vs. 0.671; two-sample t-test over *n* = 11 seeds: *p* = 1.3 *×* 10^−7^). (**D**). **Trajectory variability during** *S*_2_. Compression produces more stable trajectories (0.026 vs. 0.096; two-sample t-test over *n* = 11 seeds: *p* = 4.0 *×* 10^−10^). Error bars: bootstrap 95% CI.

We first asked whether the compressed model reproduces the intensity-dependent behavioral bias observed in rats. Psychometric curves split by *I*_2_ showed clear separation in the compressed model (4-dimensional bottleneck), with high-intensity stimuli biasing the agent toward reporting *T*_2_ longer (Figure 5B), closely matching the pattern reported by Rodrigues et al. (2024). The uncompressed model (64-dimensional bottleneck) showed weaker separation. We quantified this effect as the distance between high- and low-*I*_2_ psychometric curves at midrange durations (where *T*_1_ and *T*_2_ differ by at most 2 timesteps), since this is where the task is most ambiguous and behavioral biases are most apparent. The compressed model showed a significantly stronger bias than the uncompressed model (*p* = 0.037). Corticostriatal compression thus reproduces the key behavioral finding that stimulus intensity systematically distorts duration judgements.

We next examined whether this behavioral bias is accompanied by the same neural signatures observed in rat DLS. Both models showed intensity-modulated time codes during *S*_2_: decoded time estimates progressed faster for high-intensity stimuli, mirroring Rodrigues et al. (2024) (Figure 5C). However, the compressed model exhibited significantly stronger overall time encoding (*R*^2^ = 0.826 vs. 0.671; *p* = 1.3 *×* 10^−7^) and lower trajectory variability (0.026 vs. 0.096; *p* = 4.0 *×* 10^−10^; Figure 5D), consistent with our finding in the chunking task that corticostriatal compression produces more stable time-encoding striatal dynamics. We also reproduced the finding from Rodrigues et al. (2024) that time decoding curves during *S*_1_ collapse across intensity levels (Figure S6), consistent with *I*_1_ having little influence on temporal dynamics during the first stimulus. These results confirm that the same computational motif observed in the chunking task — stable, time-encoding striatal dynamics enhanced by compression — generalizes to a qualitatively different temporal behavior.

Our model also offers a potential reconciliation of an apparent discrepancy in the literature. The analyses of Toso et al. (2021) and Rodrigues et al. (2024) reach different conclusions from the same dataset: the former emphasizes that DLS time encoding is invariant to stimulus properties, while the latter highlights its modulation by intensity. We do not take a position on which analysis of the original data is correct, but we find that the two accounts can be reconciled theoretically by the assumptions underlying how sensory input is encoded. When we trained our model with Gaussian amplitude-coded observations rather than Poisson rate-coded observations, the compressed model reproduced the pattern reported by Toso et al. (2021): time encoding invariant to both stimulus intensity and duration (Figure S7). Under Poisson rate coding — arguably more appropriate for somatosensory spike trains driven by whisker vibrations (Arabzadeh et al., 2004; Mountcastle et al., 1969) — the same model instead produced the intensity-modulated dynamics and behavioral biases reported by Rodrigues et al. (2024). In both cases, the compressed model produced the relevant effects more strongly than the uncompressed model, suggesting that corticostriatal compression is a key mechanism regardless of input coding scheme.

#### Control of Behavior from Compressed Cortical Input

Having established that corticostriatal compression reproduces both the intensity-biased behavior and the intensity-modulated time dynamics, we next asked whether the compressed cortical representation causally drives these biases. PCA of the bottleneck activations in the compressed model revealed a clear direction separating high- and low-intensity stimuli (Figure 6A). We perturbed the bottleneck along this intensity direction during *S*_2_ presentation (Figure 6B): on trials where *I*_2_ was high, we shifted the bottleneck toward the low-intensity centroid, and vice versa. We quantified the effect using a *perturbation swing* — the summed shift in choice probability *P* (*T*_2_ *> T*_1_) across both perturbation conditions, measured in the expected direction (see Methods). The compressed model showed a significant swing while the uncompressed model did not (Figure 6C). The effect was asymmetric, with perturbation in the high-to-low direction producing a larger behavioral shift than low-to-high (Figure S8), potentially paralleling evidence that high-intensity stimuli engage somatosensory circuits more strongly (Mountcastle et al., 1969).

**Fig. 6.**
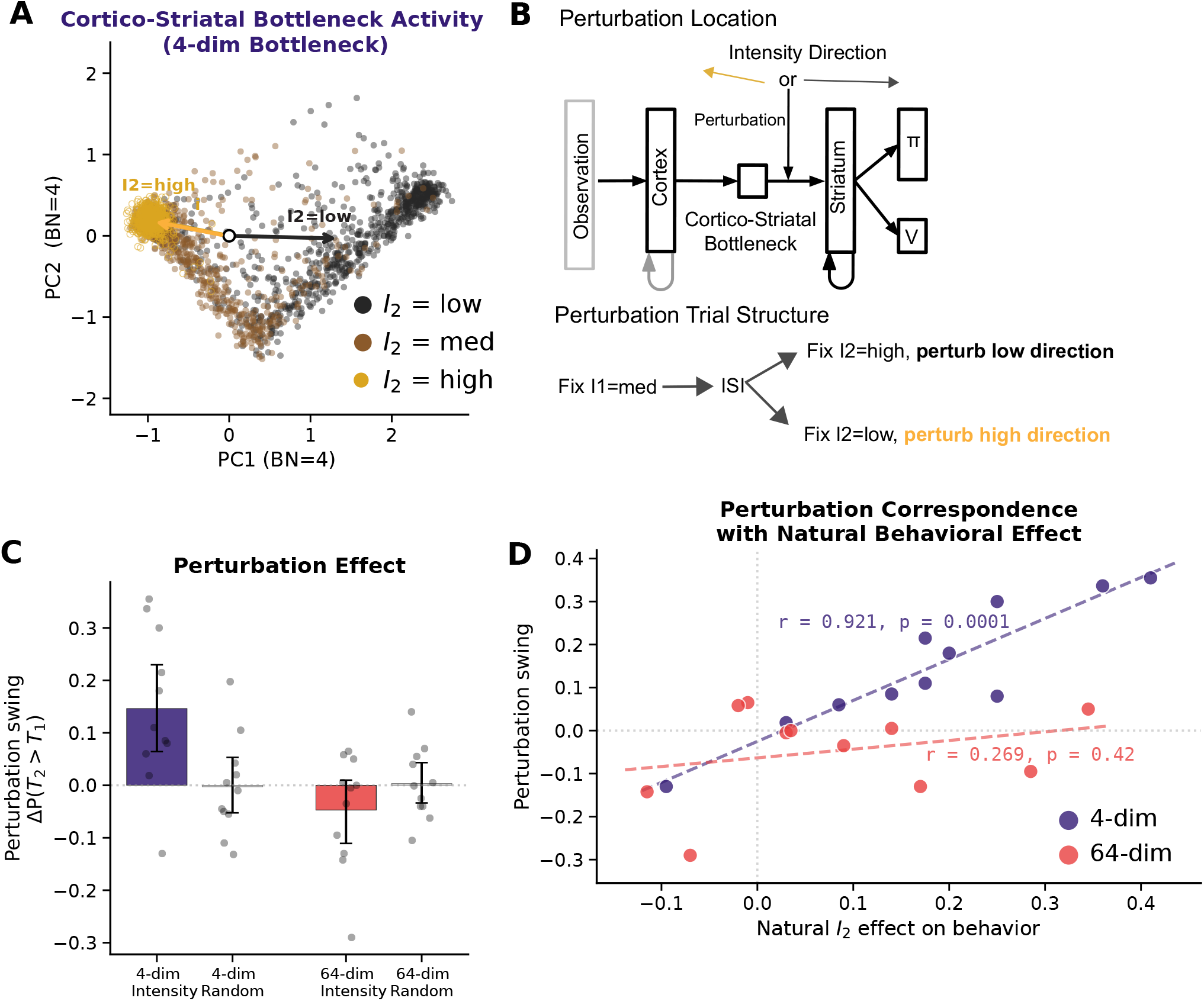
Compressed Cortical Activity Controls Intensity-Related Duration Judgement Biases. (**A**). **Bottleneck PCA**. High- and low-intensity stimuli are clearly separable in the 4-dimensional bottleneck space. (**B**). **Perturbation schematic**. With *I*_1_ fixed, the bottleneck is shifted toward the opposite intensity direction during *S*_2_: high-*I*_2_ trials are perturbed toward low, and vice versa. (**C**). **Perturbation swing** — the summed shift in *P* (*T*_2_ *> T*_1_) across both perturbation conditions, measured in the expected direction. The compressed model shows a significant swing (one-sample t-test against zero, *p* = 0.008, *n* = 11 seeds) the uncompressed model does not (one-sample t-test against zero, *p* = 0.179, *n* = 11 seeds). Perturbation along random orthogonal directions has no effect (*p* = 0.581 and *p* = 0.262 for compressed and uncompressed respectively). (**D**). **Perturbation correspondence with natural behavioral bias**. Each dot is one training seed. The compressed model shows near-perfect correspondence between natural *I*_2_ bias and perturbation susceptibility (Pearson *r* = 0.921, *p* = 0.0001); the uncompressed model shows none (Pearson *r* = 0.269, *p* = 0.42), indicating the bottleneck is the causal pathway only when capacity-limited.

The strongest evidence that the bottleneck serves as the causal pathway for intensity biases comes from the correspondence between perturbation susceptibility and natural behavioral bias. The magnitude of each seed’s perturbation swing was near-perfectly correlated with its natural *I*_2_ bias — the degree to which that seed’s unperturbed behavior was influenced by intensity (*r* = 0.921, *p* = 0.0001; Figure 6D). This is precisely what one would expect if the bottleneck is the channel through which intensity biases duration judgements: seeds whose cortical drive carries a stronger intensity signal naturally show larger behavioral biases and are most susceptible to artificial perturbation of that signal. The uncompressed model showed no such correspondence (*r* = 0.269, *p* = 0.42), indicating that without compression, intensity information is distributed across cortex and striatum and cannot be causally manipulated via a single direction. Together, these results demonstrate that corticostriatal compression consolidates task-relevant stimulus features into a compact cortical control signal that causally shapes temporal judgements through the striatum’s time-encoding dynamics.

### Motor Timing

The chunking and duration judgement tasks establish that corticostriatal compression separates cortical control from striatal dynamics across two forms of discrete decision-making. Our framework makes a further prediction: compressed cortical input should also provide low-dimensional control signals for continuous, precisely timed motor behavior. The motor timing domain offers a particularly strong test of this prediction because Hidalgo-Balbuena et al. (2019) not only recorded stable time-encoding trajectories in DLS during a well-learned motor habit, but also showed that optogenetic perturbation of somatosensory cortical input to DLS disrupts timing while preserving the structure of the habit — precisely the dissociation our framework predicts.

#### Structured Behavior and Striatal Time-Encoding Dynamics

In the task of Hidalgo-Balbuena et al. (2019), rats navigate a treadmill that moves them backward and must reach a goal zone at a precise time (Figure 7A). With practice, rats develop a stereotyped “Front-Back-Front” (FBF) strategy: they first **drift** backward passively, then **hold** position by running in place, and finally **sprint** forward to the goal. Recordings from DLS during this behavior revealed stable, time-encoding neural trajectories (Figure 7B), the kind of stereotyped temporal program that our framework predicts compression should produce.

**Fig. 7.**
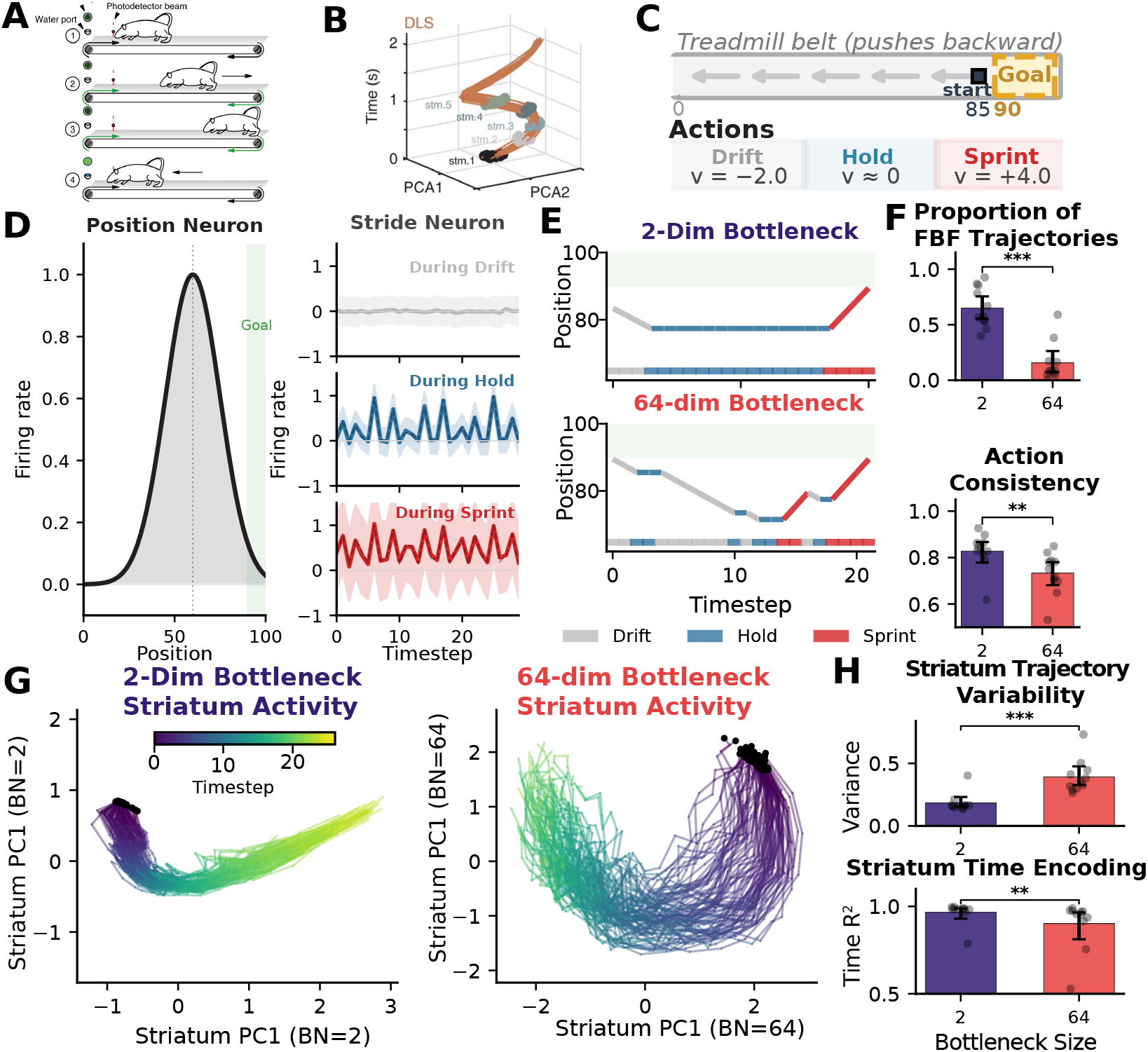
Corticostriatal Compression Produces Stereotyped Motor Timing and Stable Time-Encoding Striatal Dynamics. (**A**). **Treadmill task** of Hidalgo-Balbuena et al. (2019). Rats reach a goal zone at a specified time, developing a stereotyped Front-Back-Front (FBF) strategy. (**B**). **DLS neural trajectories**. Hidalgo-Balbuena et al. (2019) **found striatal activity organized within stable time-encoding trajectories. (C**). **Simulated task**. The model selects among three actions (drift, hold, sprint) to reach the goal zone at timestep 20. (**D**). **Observation space**. 32 position neurons (Gaussian tuning curves, left) and 32 stride neurons (periodic tuning whose clarity depends on the current action, right). (**E**). **Example action trajectories**. The compressed model (2-dimensional bottleneck) produces clean drift *→* hold *→* sprint sequences; the uncompressed model (64-dimensional bottleneck) shows disorganized switching. (**F**). **Behavioral quantification**. FBF fraction (top) and action consistency (bottom) are both significantly higher for the compressed model. (**G**). **Striatal PCA trajectories**. The compressed model produces stereotyped, temporally ordered trajectories. (**H**). **Trajectory variability and time encoding**. Variability: 0.0025 vs. 0.0198 (non-parametric two-sample Mann-Whitney *p* = 0.0001). Time encoding *R*^2^ (middle third of trial): 0.967 vs. 0.902 (non-parametric two-sample Mann-Whitney *p* = 0.009). Error bars: bootstrap 95% CI; dots are individual seeds.

We simulated this task by providing models with three discrete actions — drift (*v* = −2), hold (*v ≈* 0), and sprint (*v* = +4) — and requiring arrival at the goal zone at timestep 20 (Figure 7C). The model starts near the goal, so sprinting immediately risks arriving too early. The observation space consists of 32 position neurons with Gaussian tuning curves and 32 stride neurons with periodic tuning whose clarity depends on the current action: clean oscillations during hold, noisier during sprint, and absent during drift (Figure 7D; see Methods). This design reflects the finding of Hidalgo-Balbuena et al. (2019) that somatosensory cortical neurons carry the most temporal information during consistent, repetitive movements. Because this task is effectively two-dimensional (estimating position and time is sufficient), we used a 2-dimensional bottleneck to impose compression that is stricter than the task’s intrinsic dimensionality.

We first asked whether the compressed model spontaneously discovers the stereotyped FBF strategy observed in rats. The compressed model organized its behavior into clean drift *→* hold *→* sprint sequences, while the uncompressed model switched erratically between actions throughout each trial (Figure 7E). We quantified this with two metrics (Figure 7F). The *FBF fraction* — the proportion of trials whose action sequence, after collapsing transient blips (*≤* 2 timesteps), reduces to a sequential drift-hold-sprint ordering — was 65% for the compressed model versus 16% for the uncompressed model (non-parametric two-sample Mann-Whitney *p <* 0.001). *Action consistency* — the mean fraction of trials matching the modal action at each timestep — was also significantly higher in the compressed model (0.828 vs. 0.734, non-parametric two-sample Mann-Whitney *p* = 0.006). Corticostriatal compression thus reliably produces the stereotyped three-phase motor program observed in rats.

Does this behavioral rigidity come at the cost of task performance, or does it improve timing precision? Supplementary analyses confirmed that compression produces directed sequential flow: time-warped modal action sequences showed that every compressed-model seed converged on the FBF strategy, while uncompressed-model seeds showed heterogeneous strategies (Figure S10A,B). Conditional transition probabilities revealed a double dissociation: the compressed model showed significantly higher forward (FBF-consistent) transitions while the uncompressed model showed significantly higher backward (FBF-inconsistent) transitions (Figure S10C). Despite this rigidity, the compressed model achieved more precise motor timing: mean arrival time was significantly closer to the goal (20.4 vs. 21.8 timesteps, *p* = 0.0008), with 68% of arrived trials landing exactly at *t* = 20 compared to 43% for the uncompressed model (*p* = 0.004), while both models reached the goal at comparable rates (92%; Figure S9). The stereotyped FBF strategy is thus not merely a byproduct of compression but a functionally superior timing solution.

At the neural level, we asked whether the compressed model’s striatum shows the same stable time-encoding trajectories observed in DLS recordings. PCA of the striatal hidden states confirmed this prediction: the compressed model produced tight, temporally ordered trajectories, while the uncompressed model produced a diffuse cloud (Figure 7G). Trajectory variability was an order of magnitude lower in the compressed model (0.0025 vs. 0.0198; *p* = 0.0001; Figure 7H). To assess time encoding, we restricted the decoding analysis to the middle third of each trial, where the agent is typically in the hold phase — a stricter test than full-trial decoding, where beginning and end timesteps are trivially distinguishable by position and action context. The compressed model showed significantly stronger time encoding (*R*^2^ = 0.967 vs. *R*^2^ = 0.902, *p* = 0.009). These results confirm that the temporal scaffold produced by corticostriatal compression extends to continuous motor timing, completing the generalization across all three DLS-associated task domains.

#### Control of Behavior from Compressed Cortical Input

Our framework makes a specific prediction about cortical perturbation: if the cortex provides a low-dimensional temporal control signal while the striatum intrinsically generates the stable sequentially-ordered activity, then perturbing cortical input should shift *when* transitions occur without disrupting *what* sequence is executed. Hidalgo-Balbuena et al. (2019) tested exactly this using optogenetic perturbation of somatosensory cortical and thalamic input to DLS during the hold phase: the timing of the FBF habit shifted, but the sequential structure was preserved.

Before reproducing this perturbation, we first asked what information the corticostriatal bottleneck carries during each behavioral phase. Decoding both position and elapsed time from the bottleneck activity separately for drift, hold, and sprint timesteps (Figure 8A) revealed that in the compressed model, the bottleneck’s representational content varied markedly across actions, whereas in the uncompressed model, decoding was relatively uniform. This action-dependent information content suggests that in the compressed model, actions serve a dual role: they are not only motor commands but also gating mechanisms that control what information flows from cortex to striatum at each moment. This finding provides a principled basis for restricting our perturbation to the hold phase, when the agent relies on its internal temporal estimate to decide when to sprint.

**Fig. 8.**
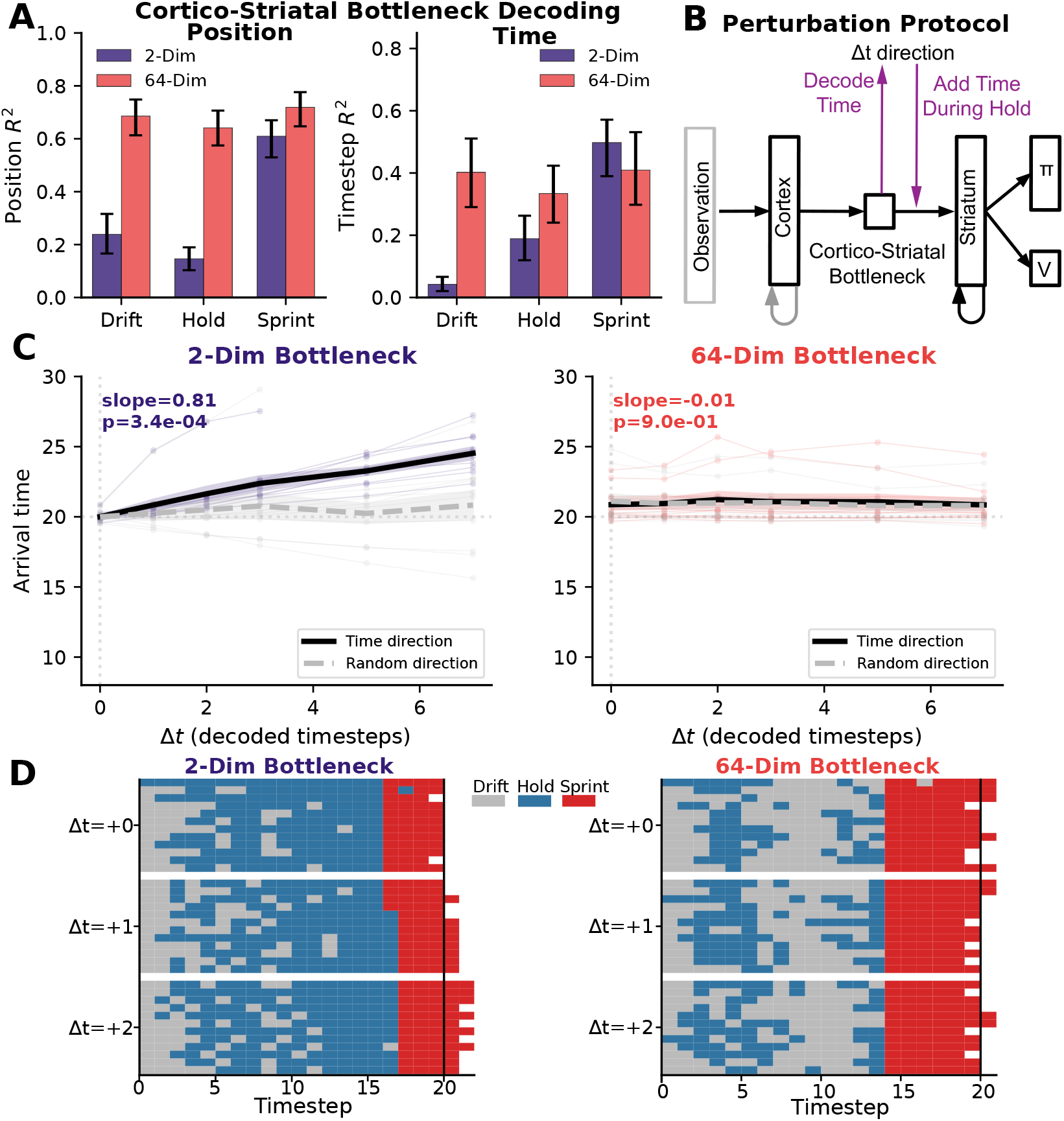
Compressed Cortical Activity Shifts Motor Timing while Preserving Sequential Structure. **(A**). Bottleneck decoding by action phase. Position (left) and time (right) decoding *R*^2^ from the bottleneck, computed separately for drift, hold, and sprint timesteps. (**B**). **Perturbation schematic**. The time-predictive direction in bottleneck space is identified via Ridge regression. Additive perturbations calibrated to shift the decoded time estimate by Δ*t* = +*k* timesteps are applied during the hold phase. (**C**). **Arrival time versus perturbation magnitude**. The compressed model shows a graded shift along the time direction (slope = 0.81, *p* = 3.4 *×* 10^−4^); random orthogonal perturbations and all uncompressed-model conditions are flat. Light lines: individual seeds; bold: mean *±* SEM (*n* = 11). (**D**). **Action rasters at** Δ*t* = 0, +1, +2. The compressed model preserves the drift *→* hold *→* sprint structure; only the hold *→* sprint transition shifts later with increasing Δ*t*. The uncompressed model lacks systematic structure.

To reproduce the optogenetic experiment of Hidalgo-Balbuena et al. (2019), we identified the direction in bottleneck space most predictive of elapsed time during the hold phase using Ridge regression and applied additive perturbations along this direction during hold only (Figure 8B). Perturbation magnitude was calibrated in units of decoded timesteps, so that Δ*t* = +*k* shifts the bottleneck’s temporal representation by the equivalent of *k* timesteps (see Methods). The compressed model showed a graded, direction-specific effect: arrival time increased linearly with perturbation magnitude along the time direction (slope = 0.81, *p* = 3.0 *×* 10^−4^), while perturbations along a random orthogonal direction had no effect (Figure 8C). The uncompressed model showed no systematic response in either direction. Critically, action rasters revealed that the compressed model preserved its drift *→* hold *→* sprint sequential structure across all perturbation levels; only the hold *→* sprint transition boundary shifted progressively later with increasing Δ*t* (Figure 8D). This directly parallels the key finding of Hidalgo-Balbuena et al. (2019): perturbation of cortical input to DLS shifts behavioral timing while leaving the motor habit intact. Together, these results demonstrate that corticostriatal compression creates a control channel in which a single direction in the cortical projection parametrically controls the timing of progression through the striatum’s behavioral program — the most direct evidence in our study for the separation of control from dynamics.

## Discussion

Across three time-sensitive behavioral paradigms associated with the dorsolateral striatum — sensorimotor chunking, duration comparison, and motor timing — we find that a single architectural constraint produces a consistent computational motif: the separation of control from dynamics between cortex and striatum. Corticostriatal compression forces the cortex to provide low-dimensional control signals while the striatum generates intrinsic, stable, time-encoding dynamics. This separation accounts for domain-specific behavioral and neural signatures — human-like chunking with action slipping, intensity-biased duration judgements with stimulus-modulated time coding, and stereotyped motor programs with perturbation-sensitive timing — and perturbation experiments confirm that the compressed cortical signal causally controls behavior while the striatum’s sequential structure remains intact. A key mechanistic insight comes from the untrained model result (Figure S4): when input dimensionality to a recurrent network is sufficiently constrained, internal dynamics dominate over input-driven variability, creating a temporal scaffold that learning can subsequently exploit (Rodrigues et al., 2024). This suggests that compression itself, not just reward-optimized compression, creates the conditions for temporal encoding. Our framework thus bridges two theoretical traditions that have developed largely in parallel: reinforcement-driven dimensionality reduction (Bar-Gad, Morris, & Bergman, 2003), which characterizes corticostriatal compression in information-theoretic terms, and the dynamical systems perspective (Haimerl et al., 2025; Lee et al., 2019; Murray & Escola, 2017), which emphasizes cortical parametric control of striatal temporal evolution. We show that these are two facets of the same mechanism: input-based compression is what creates the conditions for parametric control of stable dynamic trajectories.

The behaviors produced by our compressed models (Figure 3A, Figure 7E, Figure S10) share the defining features of habits: they are stereotyped across repetitions, inflexible to changes in sensory context, temporally rigid, and stimulus-invariant (Graybiel, 1998; Yin et al., 2004). This is consistent with the longstanding association between the DLS and habitual behavior (Yin et al., 2004), and with the proposal that chunking is a fundamental mechanism by which habits are assembled (Graybiel, 1998). Our model provides a specific account of how this arises: compression forces the cortical drive to the striatum to discard stimulus-specific detail, retaining only low-dimensional categorical signals. The striatum, receiving this impoverished but structured input, learns to generate behavior from its own internal dynamics rather than from moment-to-moment sensory guidance — a functional signature of automaticity that can also explain subcortical consolidation of automatic motor sequences (Mizes et al., 2024).

Several prior models have addressed related phenomena. Botvinick & Plaut (2004) showed that simple recurrent networks under degraded conditions produce action slips resembling human behavior, establishing that recurrence supports sequential structure; our work extends this by demonstrating that compression at the input to the recurrent network is what produces structured sequential behaviors. Nassar et al. (2018) proposed chunking as rational lossy compression in visual working memory, and Soni & Frank (2025) showed that prefrontal-basal ganglia circuits can learn adaptive chunking through dopaminergic reinforcement, with the striatum acting as a gate controller. These works complement ours by establishing the normative rationale for compression and showing how gating policies can be learned, but the basal ganglia in those models serve a switching function rather than exhibiting dynamic sequential activity. Our contribution is mechanistically distinct: we show that anatomical compression at the corticostriatal interface shapes the *geometry of striatal dynamics*, forcing the striatum into a regime where stable temporal structure dominates over input-driven variability, producing specific testable neural signatures — stable time-encoding trajectories and direction-specific cortical control that leaves striatal sequential structure intact.

Our framework relies on several simplifying assumptions that future work can relax. First, cortical recurrent weights are frozen throughout training, reflecting the assumption that cortical and striatal plasticity are governed by different learning signals operating on different timescales (Doya, 2000; Pasupathy & Miller, 2005). While this is a common modeling choice for cortical reservoirs (Buonomano & Maass, 2009; Dominey, 1995), cortical plasticity clearly exists and likely interacts with striatal learning during skill acquisition. Future work could model cortical learning explicitly and study how the co-evolution of cortical representations and striatal dynamics shapes the emergence of temporal behavior. Second, the bottleneck dimensionality in our model is hand-set rather than learned. We do not attempt to match the exact biological compression ratio; rather, the bottleneck layer is a convenient modeling device to enforce dimensionality reduction and study its functional consequences. In biology, the effective compression may be shaped by development, neuromodulation, or task demands in ways our model does not capture. Third, our comparison to neural data is indirect: we reproduce high-level behavioral and neural signatures on the same tasks used in experimental studies but do not fit neural data directly. However, our framework suggests concrete approaches for future multi-region recording analyses — for instance, measuring the effective dimensionality of one region’s projection to another and assessing the controllability of downstream dynamics from that compressed signal. Despite these limitations, the core principle — that compression at a neural interface separates control from dynamics — makes predictions that extend well beyond the specific tasks we studied.

The computational motif we describe at the corticostriatal interface also operates at the output stage of the basal ganglia. After processing within the striatum and the pallidal nuclei, basal ganglia output projects through the thalamus and back to cortex, forming the cortico-basal ganglia-thalamocortical loops that are a fundamental organizing principle of this circuit (Alexander et al., 1986). At this output stage, a second bottleneck exists: the basal ganglia output nuclei (GPi/SNr) contain far fewer neurons than their thalamic and cortical targets (Humphries, 2025). Humphries (2025) argues that the primary function of this output bottleneck is to control cortical dynamics, proposing that each output neuron sets the weight of a basis function defined by its synaptic contacts. Logiaco et al. (2021) provide a mechanistic account of how this control is implemented: basal ganglia output selects which thalamic neurons are disinhibited, and these few neurons deliver low-rank perturbations to a recurrent cortical network, controlling which motor motif is executed and when transitions occur. This is fundamentally the same principle we describe — a low-dimensional signal at the interface between regions parametrically controls dynamics within the downstream circuit. Importantly, the corticostriatal and thalamocortical bottlenecks are not independent. Haber et al. (2000) characterized their relationship as an ascending spiral: information processed through one cortico-basal ganglia loop feeds forward into the next, progressively transforming limbic signals into sensorimotor output. Our framework may thus describe a general principle that operates at multiple nodes of this spiral: compression at each interface creates the conditions for low-dimensional control of stable dynamics in the downstream circuit.

The ascending spiral architecture raises a further question: does the same compression mechanism give rise to both habitual and goal-directed behavior? An intriguing clue comes from the parallel between our architecture and the meta-reinforcement learning framework of Wang et al. (2018), which uses a similar design — a recurrent network (in their case, also an LSTM) trained by reward prediction errors that develops its own internal learning algorithm — to model goal-directed behavior typically associated with prefrontal cortex. The basal ganglia contain parallel corticostriatal loops linking different cortical regions to different striatal territories: prefrontal cortex projects to the dorsomedial striatum (DMS), associated with goal-directed behavior, while sensorimotor cortex projects to the DLS, associated with habitual behavior (Alexander et al., 1986; Haber, 2003; Yin et al., 2004). The architectural similarity between our model and Wang et al. (2018) raises the possibility that goal-directed and habitual behaviors emerge from a shared corticostriatal compression mechanism, with the distinction arising not from fundamentally different circuit computations but from differences in the cortical inputs and task demands each loop receives. Testing this hypothesis would require extending our framework to model both loops simultaneously and examining how their interaction gives rise to the well-documented transition from goal-directed to habitual control (Thorn et al., 2010).

More broadly, the principle that anatomical bottlenecks separate control from dynamics may extend to any circuit in which convergent input drives a recurrent downstream population. For example, hippocampal place cells integrate highly multimodal sensory information from diverse sources (Jeffery, 2007) and also exhibit rich temporal coding relevant for episodic memory (Eichenbaum, 2014). Recent neural network models of episodic memory that parallel hippocampus have been shown to produce stable neural trajectories which implement free recall via serial position encodings (i.e. *time codes*) that are independent of item identity (Li et al., 2025), directly paralleling the trajectories that organize action chunks in the present work (Figure 3B). Similarly, the cerebellum transforms converging, compressed sensory inputs into temporally precise neural dynamics that support conditioned responses and motor coordination (Medina et al., 2000; Wagner et al., 2019). In each case, compression at the interface between regions may jointly facilitate inter-regional control and temporal computations in the downstream circuit. If this principle proves general, it would suggest that a unified theory of inter-regional communication may be within reach — one in which the dimensionality of the projection between any two regions predicts the degree to which the downstream circuit exhibits time-encoding dynamics. Testing this prediction across brain circuits represents a natural next step for the field.

## Methods

### Model Architecture

Our model implements corticostriatal compression through three modules arranged in series (Figure 1B). The **cortical module** is an recurrent neural network layer (RNN) with 64 hidden units, whose recurrent weights are randomly initialized and *frozen* throughout training. Specifically, the recurrent-to-recurrent weights (*W*_*hh*_) and recurrent biases (*b*_*hh*_) of the encoder RNN are fixed at their initial values (PyTorch default uniform initialization, 𝒰 (−1*/*8, 1*/*8)), while all other weights remain trainable. This design reflects the assumption that cortical recurrent dynamics and striatal dynamics are learned via different learning signals and timescales.

The **corticostriatal bottleneck** is a feedforward layer that maps the cortical RNN’s output (64 dimensions) to a low-dimensional space (*d*_bn_ units), followed by additive Gaussian noise with fixed standard deviation *σ*. The noise is applied at every forward pass (both training and evaluation) and models the stochasticity of corticostriatal transmission. We used *d*_bn_ = 4 for the chunking and Toso tasks (compressing from 64 to 4 dimensions), and *d*_bn_ = 2 for the treadmill task (see task-specific sections below). The uncompressed baseline used *d*_bn_ = 64 in all tasks, effectively removing the compression constraint.

The **striatal module** is a single-layer LSTM (Hochreiter & Schmidhuber, 1997) with 64 hidden units. The LSTM receives the bottleneck output as input at each timestep and maintains its own hidden and cell states across timesteps within an episode (states are reset at episode boundaries). A linear policy head maps the LSTM hidden state to action logits (softmax policy *π*), and a separate linear value head maps to a scalar state value estimate *V*. The policy and value heads share the LSTM hidden state but use independent linear projections. All striatal weights (LSTM parameters, policy head, value head) and corticostriatal weights (bottleneck projection, encoder input weights) are trained via reinforcement learning. Cortical recurrent weights remain frozen.

### Training Algorithm

All models were trained using Proximal Policy Optimization (PPO; Schulman et al., 2017), a policy gradient algorithm that stabilizes training by clipping the policy update ratio. The PPO objective is:

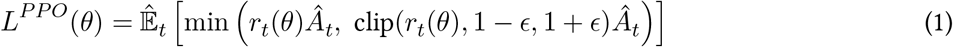

where 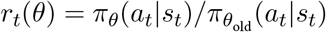 is the probability ratio between the current and previous policy, *Â*_*t*_ is the estimated advantage, and *ϵ* = 0.2 (the RLlib default). Parameters were optimized using Adam (Kingma & Ba, 2014). The LSTM hidden and cell states were initialized to zero at the beginning of each episode.

Training was implemented using RLlib (Liang et al., 2018). For each task and bottleneck condition, we trained 11 independent models with seeds {0, 10, 20, …, 100} to ensure robustness to parameter initialization and environment stochasticity. Default hyperparameters are listed in Table 1; task-specific deviations are noted in each task section.

**Table 1:** Default PPO training hyperparameters. Task-specific deviations noted in text.

| Hyperparameter | Default Value |
| --- | --- |
| Learning rate | $1 \times 10^{-3}$ |
| Batch size | 128 |
| Discount factor $\gamma$ | 0.99 |
| Bottleneck noise $\sigma_{\text{bn}}$ | 0.4 |

### Action Chunking Task

We used a serial decision-making task inspired by Lai et al. (2025). At each timestep *t ≤ T*, a stimulus *s*_*t*_ is drawn from a set *S* = {0, 1, …, 15} of 16 possible stimuli. Stimuli are presented as one-hot vectors. Each stimulus *s*_*i*_ has a single correct action *a*_*i*_ = *i*. The agent receives a reward of 0 for correct actions and a penalty of −1*/T* for incorrect actions.

A subset *F* = {0, 1, …, 7} of 8 stimuli forms a “chunk” that frequently appears in a fixed temporal order. At each timestep not already occupied by the chunk sequence, there is a probability *P*_init_ = 0.45 that the chunk sequence initiates (provided the remaining episode length can accommodate the full sequence). When initiated, chunk stimuli are presented in order across consecutive timesteps. When the chunk is not initiated, each stimulus is drawn randomly: chunk stimuli appear with probability *P*_aux_ = 0.09 each, and non-chunk stimuli fill the remaining probability mass. Episode length was fixed at *T* = 8 timesteps.

The observation at each timestep consisted of the stimulus one-hot vector (16 dimensions), a one-hot encoding of the previous action (16 dimensions), and the previous reward (1 dimension), yielding a 33-dimensional observation consistent with the meta-RL framework (Wang et al., 2018) that helps models learn fast adaptation within trials. Models were trained for 3,000 epochs with a learning rate schedule decaying from 1 *×* 10^−3^ to 2 *×* 10^−4^ at epoch 2,800. Bottleneck noise was *σ*_bn_ = 0.4.

For the IID control environment, the same trained models were evaluated on episodes where all 16 stimuli were drawn independently with equal probability (no chunk structure). This tests whether time-encoding properties persist in the absence of temporal regularities.

### Poisson Duration Comparison Task

We simulated the duration comparison task from Toso et al. (2021), with observations designed to capture the statistical properties of somatosensory spike trains as highlighted by the reanalysis of Rodrigues et al. (2024).

#### Trial structure

Each trial consisted of five phases presented sequentially: a pre-stimulus period (2 timesteps), stimulus *S*_1_ (*T*_1_ timesteps), an inter-stimulus interval (ISI; 16 timesteps), stimulus *S*_2_ (*T*_2_ timesteps), and a post-stimulus period (2 timesteps), followed by a single response timestep. Stimulus durations *T*_1_ and *T*_2_ were independently sampled as integers from {2, 3, …, 8} with the constraint *T*_1_ ≠ *T*_2_. Stimulus intensities *I*_1_ and *I*_2_ were independently sampled from three levels: low (*λ* = 0.6), medium (*λ* = 1.2), and high (*λ* = 2.4), expressed as Poisson rate parameters.

#### Observations

At each timestep, the agent observed a 16-dimensional vector of spike counts drawn from independent Poisson distributions. During stimulus periods, each of the 16 “cochlear” neurons fired with rate equal to the current stimulus intensity (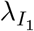 during *S*_1_, 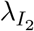 during *S*_2_). During non-stimulus periods (pre-stimulus, ISI, post-stimulus), neurons fired at a baseline rate of *λ*_base_ = 0.42. This observation model entangles intensity and duration: the total spike count during a stimulus is approximately 16 *× λ × T*, making it impossible to perfectly separate intensity from duration based on total activity alone. The observation was augmented with a one-hot encoding of the previous action (2 dimensions) and the previous reward (1 dimension), yielding a 19-dimensional input.

#### Task and reward

On the response timestep, the agent selected one of two actions: 0 (“*T*_1_ *> T*_2_”) or 1 (“*T*_2_ *> T*_1_”). Correct responses received a reward of 1.0; incorrect responses received 0.0. No reward was given during non-response timesteps.

#### Training

Models were trained for 1,000 epochs with a learning rate decaying from 5 *×* 10^−4^ to 5 *×* 10^−5^, a batch size of 1,024, 5 parallel environments per worker, and bottleneck noise *σ*_bn_ = 0.1.

#### Gaussian variant (supplementary)

For comparison with the original analysis of Toso et al. (2021), we also trained models on a Gaussian-observation variant of the task. In this version, each stimulus was represented as a 16-dimensional vector with 5 signal dimensions (mean = *I, σ* = 0.05) and 11 noise dimensions (mean = 0, *σ* = 0.1), shuffled randomly at each timestep to prevent position-specific encoding. Intensities were drawn from {0.3, 0.6, 0.9}. The ISI was 4 timesteps and the bottleneck noise was *σ*_bn_ = 0.4. Models were trained for 250 epochs.

### Treadmill Motor Timing Task

We developed a reinforcement learning environment inspired by the treadmill navigation task of Hidalgo-Balbuena et al. (2019).

#### Environment

The agent navigates a one-dimensional track of length 100 units. A virtual treadmill displaces the agent backward at a constant speed of 2.0 units per timestep. The agent starts at a position drawn uniformly from [75, 89.5] (i.e., 85.0 *±* 10.0, clipped to remain below the goal zone). The goal zone begins at position *x* = 90.0. The agent must reach the goal zone at or after timestep 20 (the target arrival time); arriving before timestep 20 incurs a harsh penalty. The maximum episode duration is 30 timesteps.

#### Actions

The agent selects from three discrete actions at each timestep:

- **Drift** (action 0): no self-generated motion; the treadmill pushes the agent backward at −2.0 units/step.
- **Hold** (action 1): the agent runs at the treadmill speed, producing net velocity *≈* 0.
- **Sprint** (action 2): the agent runs at maximum speed (6.0 units/step), producing net forward velocity of +4.0 units/step.

These actions correspond to the three phases of the rat FBF strategy observed by Hidalgo-Balbuena et al. (2019): passive backward drift, steady-state running in place, and forward sprinting to the goal.

#### Observation space

The observation consisted of two populations of simulated cortical neurons (64 neurons total), designed to capture the key statistical properties of somatosensory input during treadmill locomotion:

##### Position neurons (*n* = 32)

Gaussian tuning curves tiled uniformly across the track. Neuron *i* with preferred position *p*_*i*_ fires with rate:

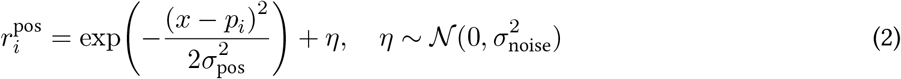

where *x* is the agent’s current position, *σ*_pos_ = 15.0, and *σ*_noise_ = 0.3.

##### Stride neurons (*n* = 32)

von Mises (circular Gaussian) tuning curves encoding the phase of a periodic stride signal. The stride phase advances at a fixed frequency of *f*_stride_ = 0.37 cycles per timestep (*ϕ*_*t*_ = 2*πf*_stride_ *· t*). Neuron *j* with preferred phase *θ*_*j*_ fires with rate:

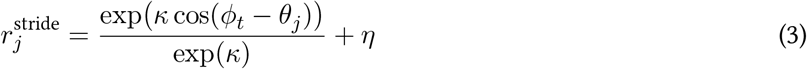

Critically, the quality of this signal depends on the agent’s current action:

- During **hold**: clean signal with concentration *κ* = 3.0 and noise *σ* = 0.3, reflecting rhythmic somatosensory feedback from steady locomotion.
- During **sprint**: degraded signal with *κ* = 1.0 and noise *σ* = 0.8, reflecting noisier proprioceptive input during acceleration.
- During **drift**: pure noise (*σ* = 0.3, no von Mises signal), reflecting the absence of self-generated movement.

This design is motivated by the finding of Hidalgo-Balbuena et al. (2019) that somatosensory cortical neurons carry the most temporal information during consistent, rhythmic movement phases (i.e., the hold phase of the FBF strategy). Because the treadmill task specifically tests how cortical sensory input serves as a temporal reference for the striatum (Hidalgo-Balbuena et al., 2019), we provided action information only through the sensory observation itself — via action-dependent stride neuron tuning — rather than as an explicit label, so that the temporal reference signal and the action feedback share the same noisy, capacity-limited channel through the bottleneck. Unlike the other two tasks, action information is already embedded in the sensory input via the action-dependent phase neurons and and thus does not need to be provided as a separate channel.

#### Reward function

Reward is given to the agent at the end of the trial. The reward was designed to encourage precise timing of goal arrival:

If the agent entered the goal zone (*x ≥* 90.0) before timestep 20: a penalty of −1.0 and the episode terminated.

- If the agent entered the goal zone at timestep *t ≥* 20: a reward of exp(−0.3 *·* (*t* − 20)) and the episode terminated. This exponentially decaying reward strongly favors on-time arrival (reward = 1.0 at *t* = 20, *≈* 0.74 at *t* = 21, *≈* 0.55 at *t* = 22).
- If the agent did not enter the goal zone by timestep 30: a timeout penalty of −0.2.

#### Training

Models were trained for 125 epochs with a learning rate decaying from 3 *×* 10^−4^ to 5 *×* 10^−5^, a batch size of 1,024, an entropy coefficient of 0.05 (to encourage exploration of the three-action space), and bottleneck noise *σ*_bn_ = 0.1. Because the task is effectively two-dimensional (the agent must estimate its position and elapsed time to determine when to sprint), we used a 2-dimensional bottleneck to impose compression that is lower than the dimensionality of the task (note that, with the added noise to the bottleneck, the model will not be able to perfectly represent both position and time).

### Trajectory Analysis

We computed two primary metrics on the full 64-dimensional LSTM hidden state to characterize the geometry of striatal neural trajectories. These metrics were applied identically across all three tasks.

#### Trajectory variance

For each seed and condition, we collected hidden state trajectories across multiple trials. At each timestep *t*, we computed the variance of each hidden state dimension across trials, then averaged across dimensions:

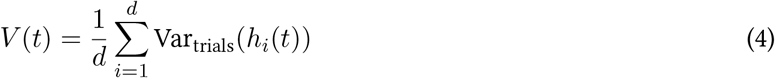

where *h*_*i*_(*t*) is the *i*-th dimension of the hidden state at timestep *t* and *d* = 64. Trajectory variance is then the mean of *V* (*t*) across timesteps. Lower values indicate more stereotyped, trial-invariant dynamics.

#### Time encoding

We trained a Ridge regression model to predict elapsed timestep from the full hidden state vector. Performance was evaluated using 5-fold cross-validation. The metric is the mean cross-validated *R*^2^. For the treadmill task, we additionally restricted the decoding analysis to the middle third of each trial to provide a stricter test: the beginning (drift) and end (sprint) phases are trivially distinguishable by position and action context, so mid-trial decoding is a stricter measure of temporal representations.

### Perturbation Experiments

Across all three tasks, we employed a common perturbation framework to test whether compressed cortical representations causally control behavior. The general procedure involves three steps: (1) identify a task-relevant direction in the bottleneck space, (2) apply additive perturbations along this direction during task execution, and (3) measure the behavioral effect relative to a random orthogonal control direction.

#### General framework

Perturbations were applied via a forward hook registered on the Gaussian noise layer of the bottleneck. At each forward pass, the hook added a perturbation vector *δ* = *α · ŵ* to the bottleneck output, where *ŵ* is the unit-length direction of interest and *α* is the perturbation magnitude. The random control direction was generated by drawing a random vector and orthogonalizing it against *ŵ* via Gram-Schmidt, then normalizing. The random control used the same bottleneck-space magnitude *α*, ensuring a fair comparison.

#### Chunking perturbation

The chunk direction was identified from PCA of the bottleneck activations. We computed the centroid of bottleneck states during chunk stimuli and non-chunk stimuli separately, and defined the chunk direction as the vector from the non-chunk centroid to the chunk centroid. The empirical magnitude was defined as the norm of this vector. Perturbations were applied during random (IID) episodes at every timestep, with magnitudes varied parametrically from 0 to 3.0 in units of the empirical magnitude. The behavioral metric was the number of correctly timed chunk actions per episode: the count of timesteps where the model produced the action that would have been correct if the chunk sequence were being presented at that position.

#### Toso perturbation

The intensity direction was identified from PCA of the bottleneck activations during *S*_2_. We computed the centroid of bottleneck states for high-*I*_2_ and low-*I*_2_ trials separately, and defined the intensity direction as the vector from low to high. Perturbations were applied during *S*_2_ only, in a counterbalanced design: on trials with *I*_2_ = high, the bottleneck was shifted toward the low-intensity centroid (magnitude = −1.0*×* empirical magnitude), and on trials with *I*_2_ = low, the bottleneck was shifted toward the high-intensity centroid (magnitude = +1.0). The perturbation swing was defined as the summed shift in choice probability *P* (*T*_2_ *> T*_1_) across both conditions, measured in the expected direction:

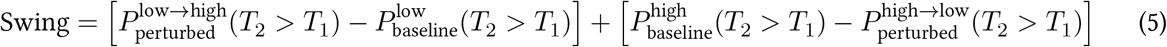

A positive swing indicates that the perturbation shifted behavior in the expected direction for both conditions.

#### Treadmill perturbation

The time direction was identified via Ridge regression from bottleneck activations to elapsed timestep fit on hold-phase data. The direction was defined as the normalized weight vector *ŵ* = *w/* ∥*w*∥, where *w* is the Ridge regression coefficient vector. Perturbation magnitude was calibrated in units of decoded timesteps: adding *ŵ ·* (Δ*t/* ∥*w*∥) to the bottleneck output shifts the linear decoder’s prediction by exactly Δ*t* timesteps. Perturbation was applied only during the hold phase (action = 1), when the agent is using its internal temporal estimate to time the hold→sprint transition. This mirrors the experimental design of Hidalgo-Balbuena et al. (2019), who optogenetically perturbed cortical input during the sustained running phase. Magnitudes were varied from Δ*t* = 0 to Δ*t* = +7 decoded timesteps. The behavioral metrics were arrival time (the first timestep at which the agent’s position exceeded the goal zone), sprint onset (the first timestep with action = 2), and per-seed regression slope of arrival time versus Δ*t*.

### Bottleneck Decoding by Action Phase (Treadmill)

To characterize what information the corticostriatal bottleneck carries during different behavioral phases, we decoded position and elapsed time from the bottleneck activations using Ridge regression with 5-fold cross-validation. Decoding was performed separately on timesteps corresponding to each action (drift, hold, sprint), yielding phase-specific *R*^2^ values for both position and time.

### Behavioral Metrics (Treadmill)

#### FBF fraction

For each trial, the action sequence was simplified using run-length encoding: consecutive timesteps with the same action were collapsed into a single block, and transient blocks (*≤*2 timesteps) were merged into the adjacent block. A trial was classified as FBF-consistent if the simplified sequence consisted of a sequential ordering containing drift, hold, and sprint phases (e.g., [0, 1, 2], [0, 2], or [1, 2]). The FBF fraction is the proportion of trials classified as FBF-consistent.

#### Action consistency

At each timestep *t*, we identified the modal (most frequent) action across all trials for a given seed. Action consistency is the mean fraction of trials matching the mode, averaged across timesteps. This metric captures how stereotyped the behavioral policy is across repeated executions, including across trials with different starting positions (75–89.5).

## Supporting information

**Fig. S1.**
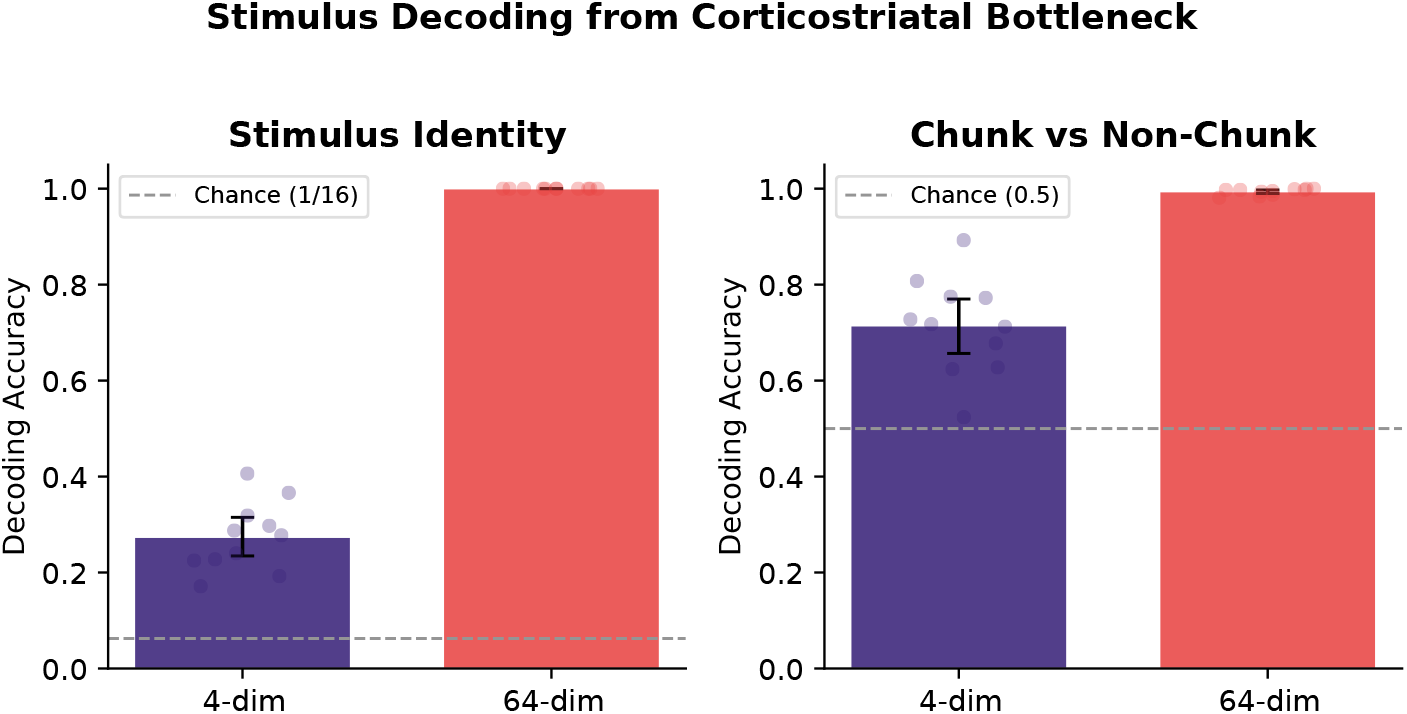
Stimulus information is lost through the corticostriatal bottleneck. Decoding accuracy from the bottleneck representation for stimulus identity (**left**) and chunk membership (**right**). Dots show individual seeds (*n* = 11); bars and error bars show means with 95% bootstrap CIs. Dashed lines indicate chance level. **Left:** A linear classifier trained to decode the current stimulus identity (16-way classification) achieves only 27% accuracy from the 4-dim bottleneck compared to 100% from the 64-dim bottleneck (*p <* 0.0001), confirming that individual stimulus identity is largely lost through compression. **Right:** A binary classifier for chunk membership (chunk vs. non-chunk stimulus) achieves 71% from the 4-dim bottleneck versus 99% from the 64-dim bottleneck (*p <* 0.0001). Although reduced, chunk membership information is partially preserved, consistent with the bottleneck retaining categorical rather than stimulus-specific information.

**Fig. S2.**
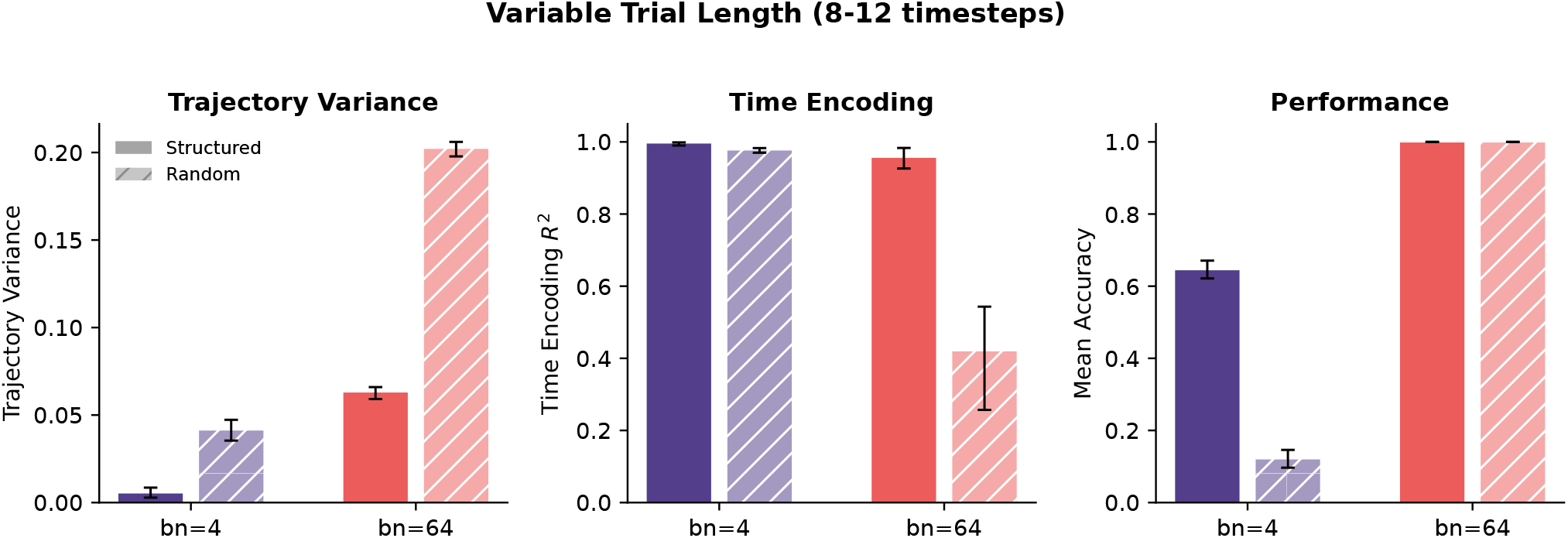
Core sensorimotor chunking results replicate with variable trial lengths. Models trained with trial lengths drawn uniformly from 8–12 timesteps (so the chunk can start anywhere from timestep *t* = 0 to *t* = 3). Single seed; error bars show 95% bootstrap CIs across episodes or cross-validation folds. **Left:** Trajectory variance remains lower for the compressed model in both structured (solid) and random (hatched) trials. **Middle:** Time encoding *R*^2^ remains near ceiling (~0.99) for the compressed model, confirming that the striatal hidden state tracks absolute elapsed time even when trial duration is unpredictable. **Right:** Chunking effect. The structured trial advantage persists: compressed models perform significantly better on structured than random trials, while uncompressed models are at ceiling for both.

**Fig. S3.**
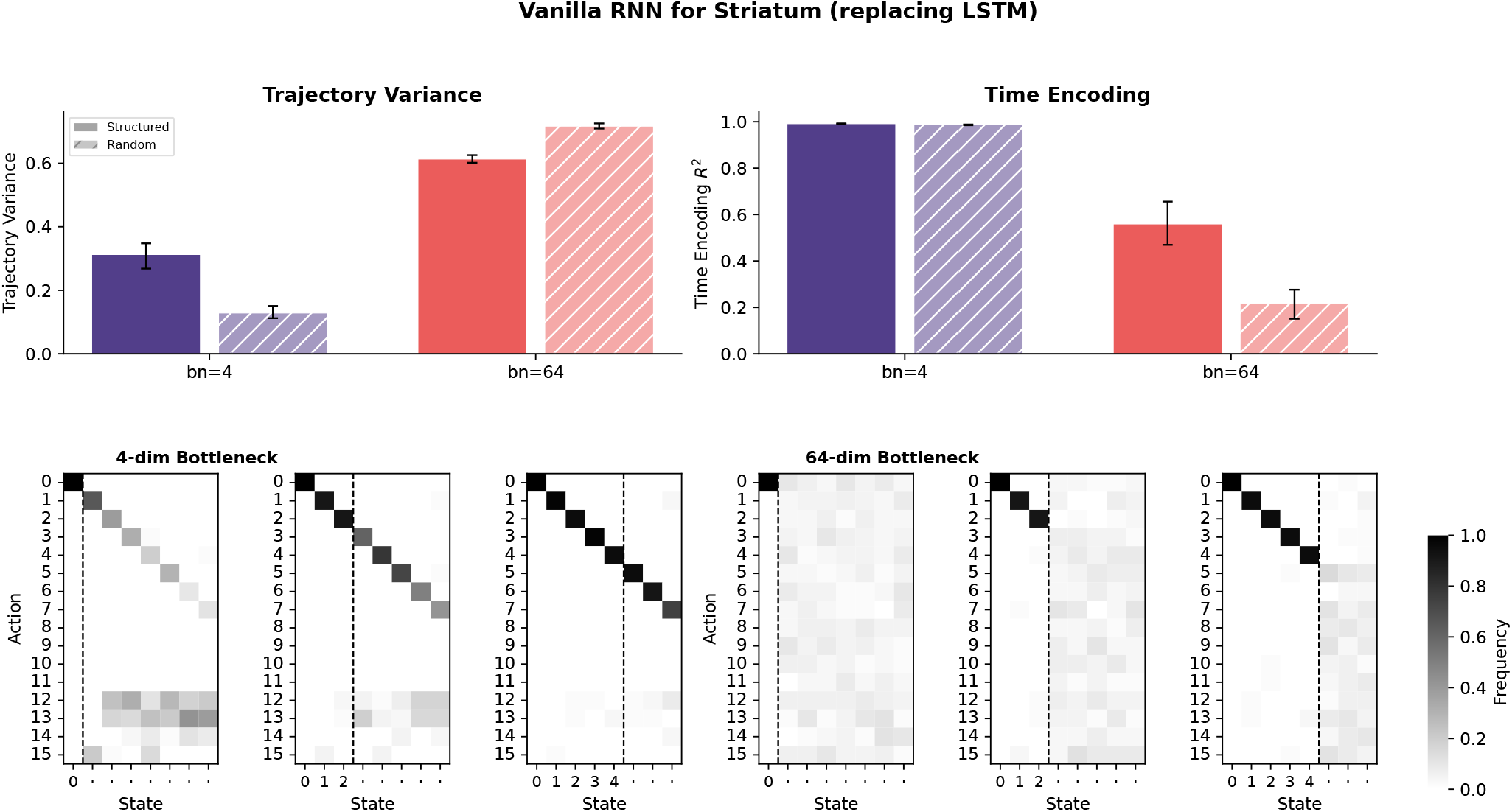
Core sensorimotor chunking results replicate with a vanilla RNN replacing the LSTM. The LSTM module in the striatal network was replaced with a vanilla RNN to test whether gating mechanisms are necessary for compression-induced chunking. **Top left:** Trajectory variance is lower for the compressed model (bn=4) than the uncompressed model (bn=64), replicating the LSTM result. **Top right:** Time encoding *R*^2^ is near ceiling for bn=4 and substantially lower for bn=64, again replicating the LSTM finding. **Bottom:** Action slipping heatmaps for three chunk subset sizes. The compressed model (**left three panels**) shows the characteristic diagonal extension past the dashed line, indicating action slipping beyond the chunk boundary. The uncompressed model (**right three panels**) shows diffuse action distributions with no slipping.

**Fig. S4.**
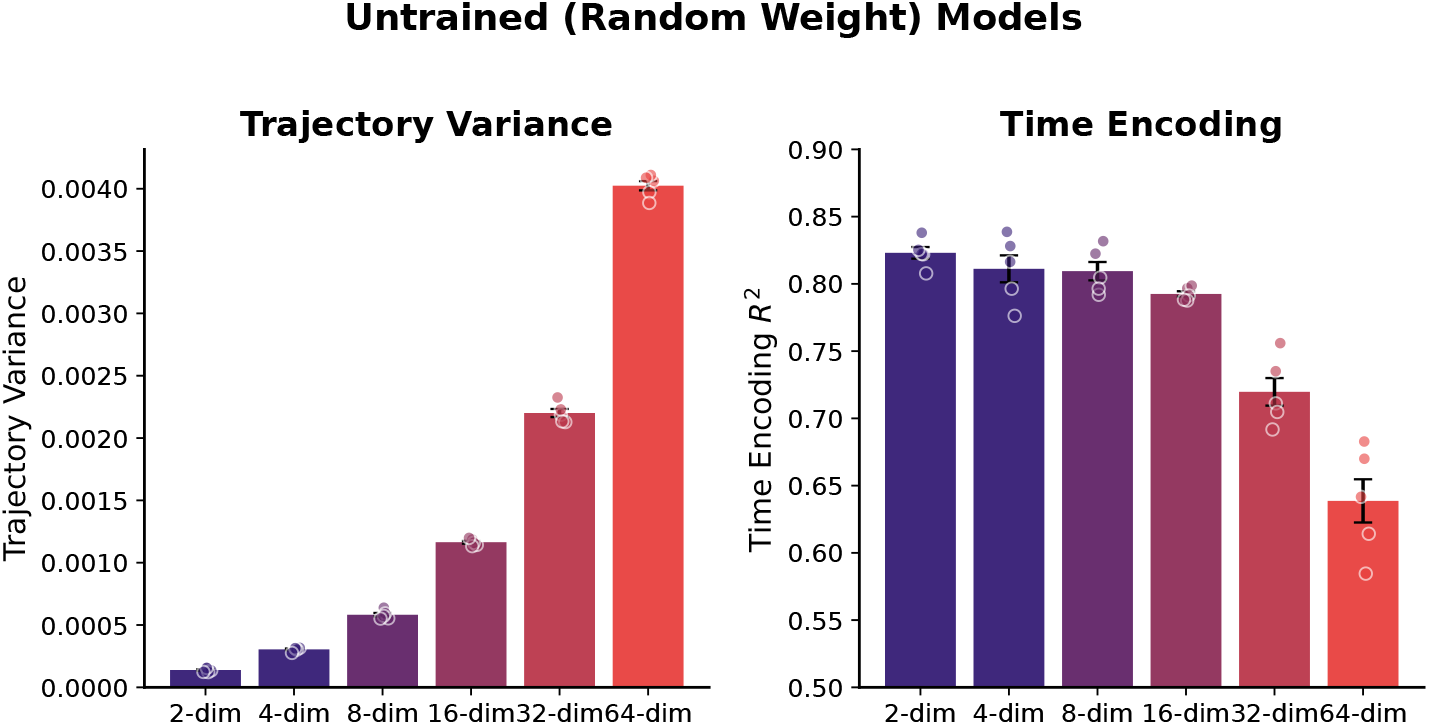
Compression creates temporal structure prior to learning. Trajectory variance (**left**) and time encoding *R*^2^ (**right**) for models with completely random (untrained) weights, evaluated on IID stimuli. We show models with different cortico-bottleneck sizes: 2-dim, 4-dim, 8-dim, 16-dim, 32-dim, and 64-dim. Dots show 5 random initializations; bars show means with 95% bootstrap CIs. More compression in the cortico-striatal bottleneck layer leads to less trajectory variance and stronger time encoding.

**Fig. S5.**
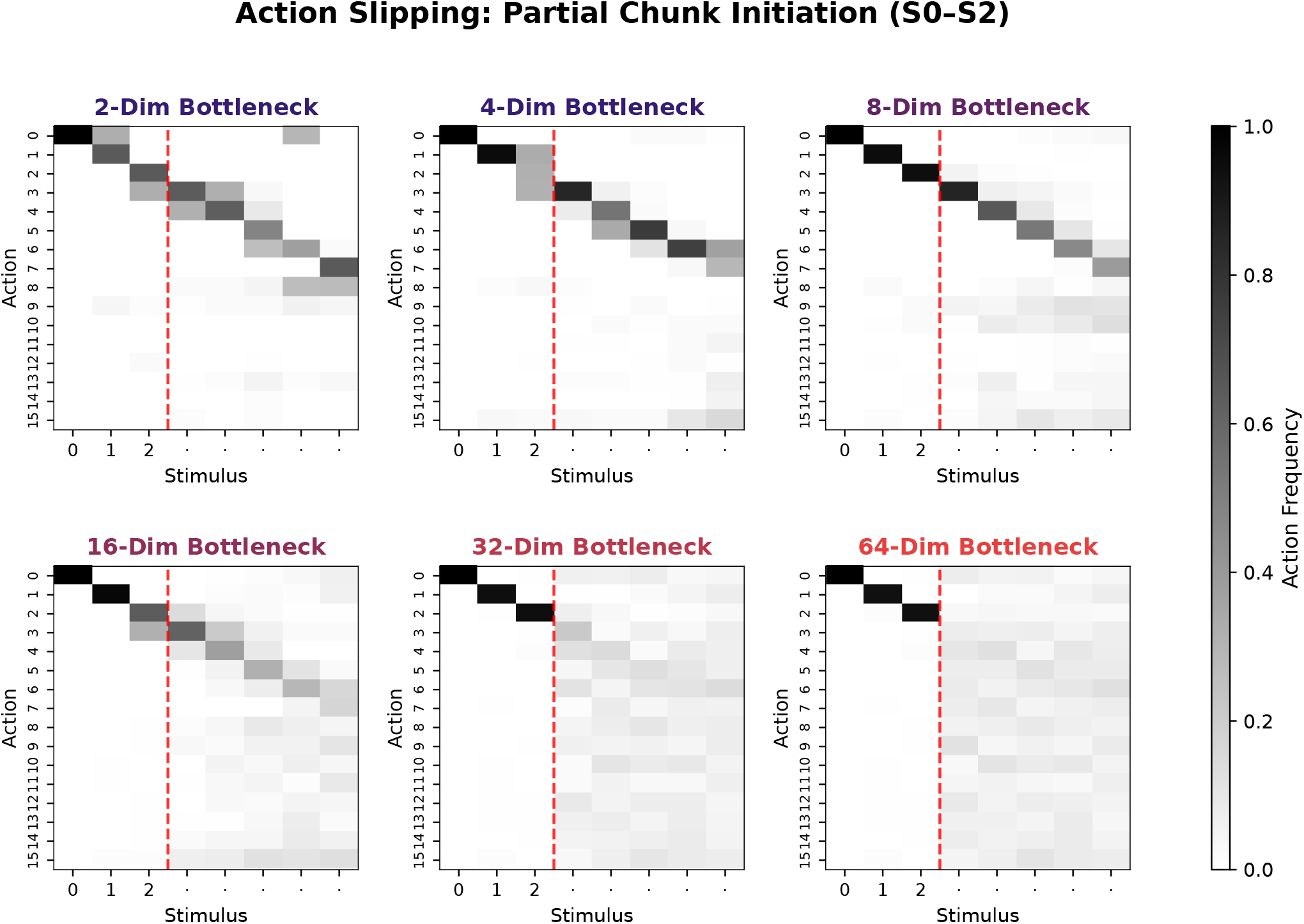
Action slipping across bottleneck dimensions. Action frequency heatmaps following partial chunk initiation (stimuli S0–S2 presented in chunk order, followed by random non-chunk stimuli). Action slipping is strongest in the most compressed models and slowly diminishes as the dimensionality of the bottleneck layer increases, with 16 dimensions (the number of stimuli in the task) being a transition point. This demonstrates a graded relationship between corticostriatal compression and the inflexibility of learned action sequences. Each panel shows the mean action frequency averaged across 3 training seeds. The 4-dim panel here may differ visually from the corresponding panel in Figure 3A, which averages across 11 seeds, but the qualitative pattern is consistent.

**Fig. S6.**
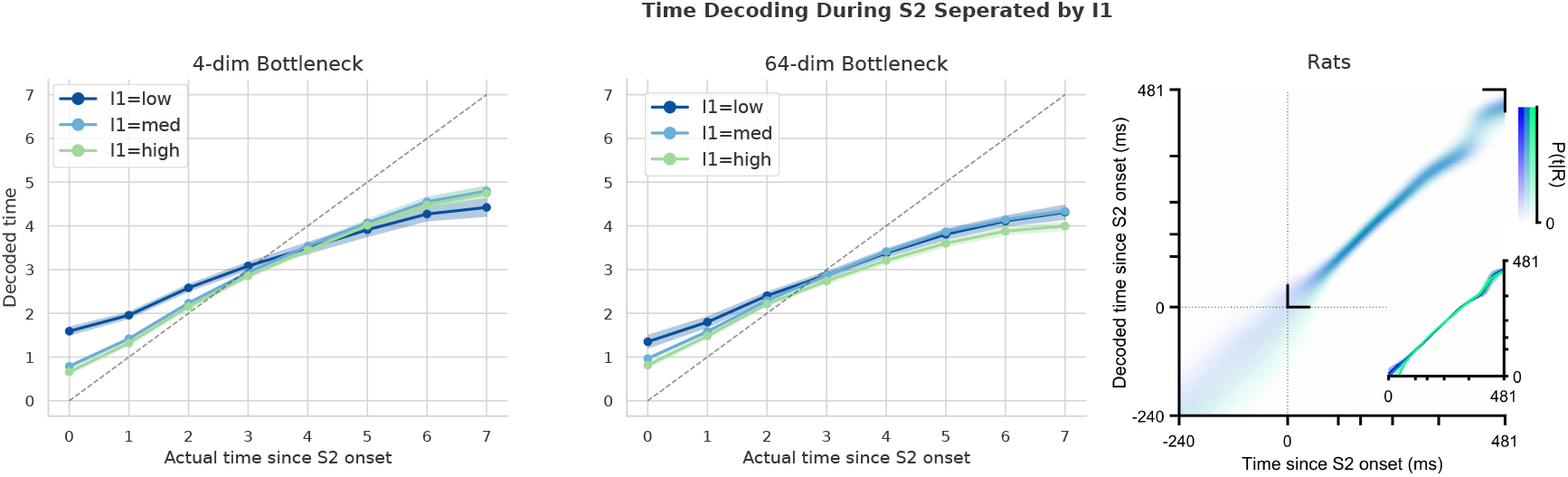
Time decoding during S2 is largely invariant to I1. Time decoding curves during the second stimulus period (S2), split by I_1_ intensity (low, medium, high), for the compressed (**left**, bn=4) and uncompressed (**right**, bn=64) models trained on the Poisson duration comparison task. Lines show cross-validated ridge regression predictions averaged across seeds; shaded regions indicate SEM. In both models, the three intensity curves largely overlap, indicating that I_1_ has minimal influence on temporal dynamics during S1. This replicates the same finding of Rodrigues et al. (2024) (shown on the far right).

**Fig. S7.**
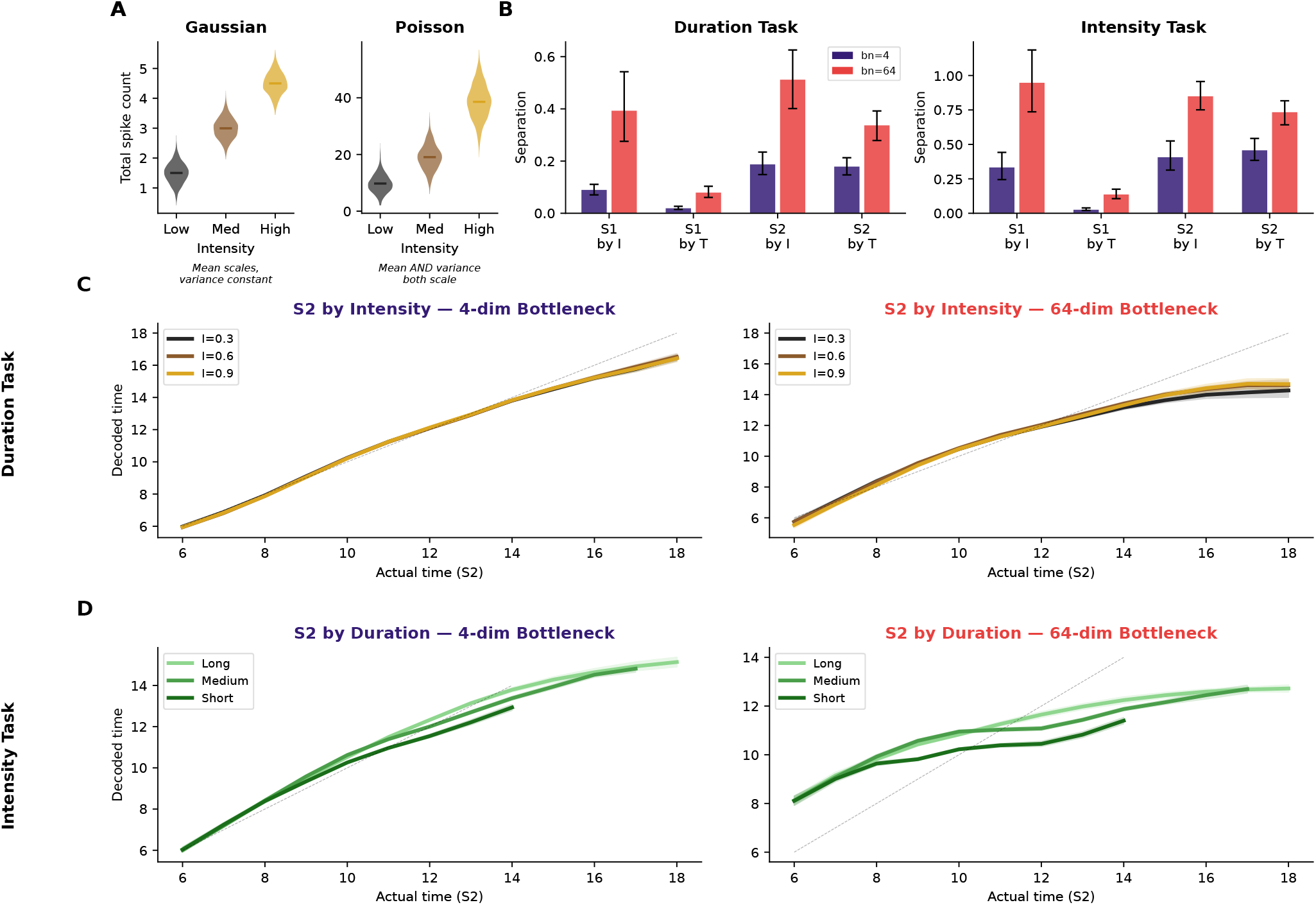
Gaussian observations produce stimulus-invariant time coding, reconciling Toso and Rodrigues findings. **(A)** Observation statistics for Gaussian (left) and Poisson (right) models. Violin plots show the distribution of total spike counts across neurons at three intensity levels. In the Gaussian model, only the mean scales with intensity while variance remains constant. In the Poisson model, both mean and variance scale with intensity, creating intensity-dependent temporal dynamics. **(B)** Mean separation of time decoding curves across all conditions (S1/S2 *×* intensity/duration splits) for the Gaussian duration task (left) and intensity task (right). Purple: bn=4; red: bn=64. The compressed model consistently shows lower separation across all conditions (all *p <* 0.001), indicating more stimulus-invariant time coding. **(C)** Example time decoding curves from the Gaussian duration task: S2 split by I_2_ intensity for bn=4 (left) and bn=64 (right). The three intensity curves nearly overlap for the compressed model, matching the stimulus-invariant pattern reported by Toso et al. (2021). **(D)** Example curves from the Gaussian intensity task: S2 split by duration for bn=4 (left) and bn=64 (right), showing the same pattern. These results demonstrate that the same compression architecture produces Toso-like invariance with Gaussian observations and Rodrigues-like modulation with Poisson observations.

**Fig. S8.**
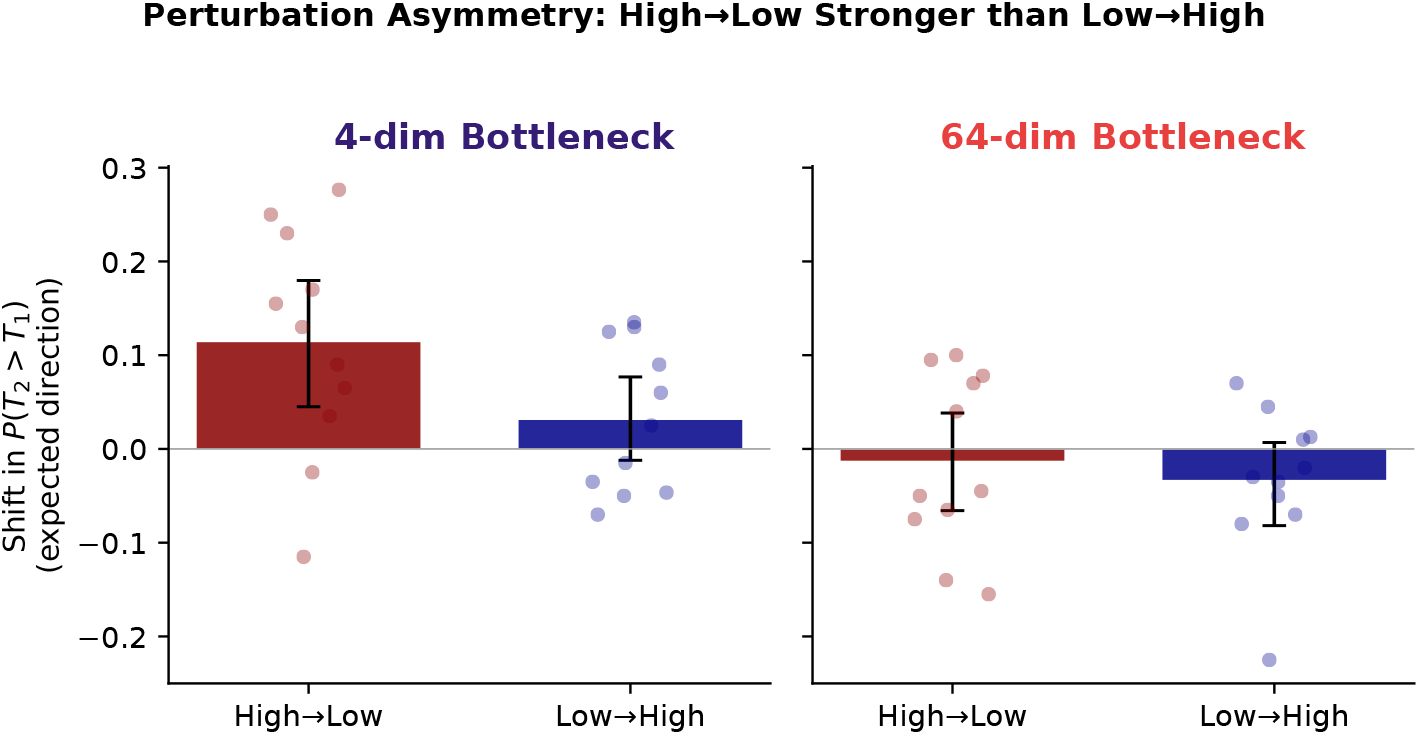
Perturbation effects in duration comparison are asymmetric: high-to-low shifts are stronger than low-to-high. Behavioral shift in *P* (*T*_2_ *> T*_1_) following bottleneck perturbation in the task-relevant direction (I_2_), separated by perturbation direction. Bars show means across 11 seeds; dots show individual seeds; error bars show 95% bootstrap CIs. **Left (bn=4):** Perturbing the bottleneck representation from high toward low I_2_ produces a significantly larger behavioral shift than the reverse direction (|high *→* low| = 0.14 vs. |low *→* high| = 0.07; paired *t*-test, *p* = 0.036). **Right (bn=64):** The uncompressed model shows no significant perturbation effect in either direction (*p* = 0.33), as expected given that perturbations to a 64-dimensional representation produce negligible changes relative to the full representational capacity.

**Fig. S9.**
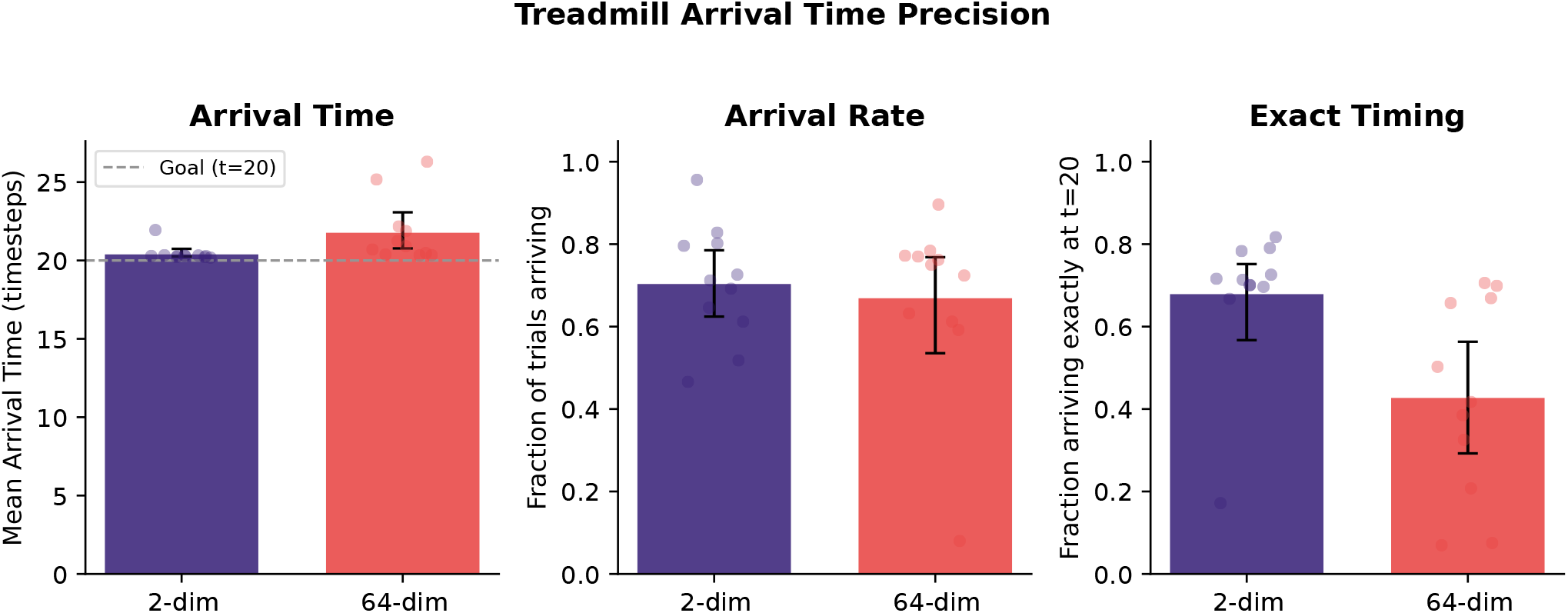
Compressed models achieve more precise motor timing. Arrival time precision on the treadmill task for compressed (bn=2, purple) and uncompressed (bn=64, red) models. Dots show individual seeds (*n* = 11); bars show means with 95% bootstrap CIs. **Left:** Mean arrival time. The compressed model arrives significantly closer to the goal time of *t* = 20 (20.4 vs. 21.8 timesteps; Mann–Whitney *U, p* = 0.0008). Dashed line indicates the goal. **Middle:** Fraction of trials in which the agent reaches the goal zone. Both models arrive at comparable rates (70.5% vs. 67.0%; *p* = 0.92), indicating that the two strategies are equally successful at reaching the goal. **Right:** Fraction of arrived trials where the agent arrives exactly at *t* = 20. The compressed model is significantly more precise (68% vs. 43%; *p* = 0.004).

**Fig. S10.**
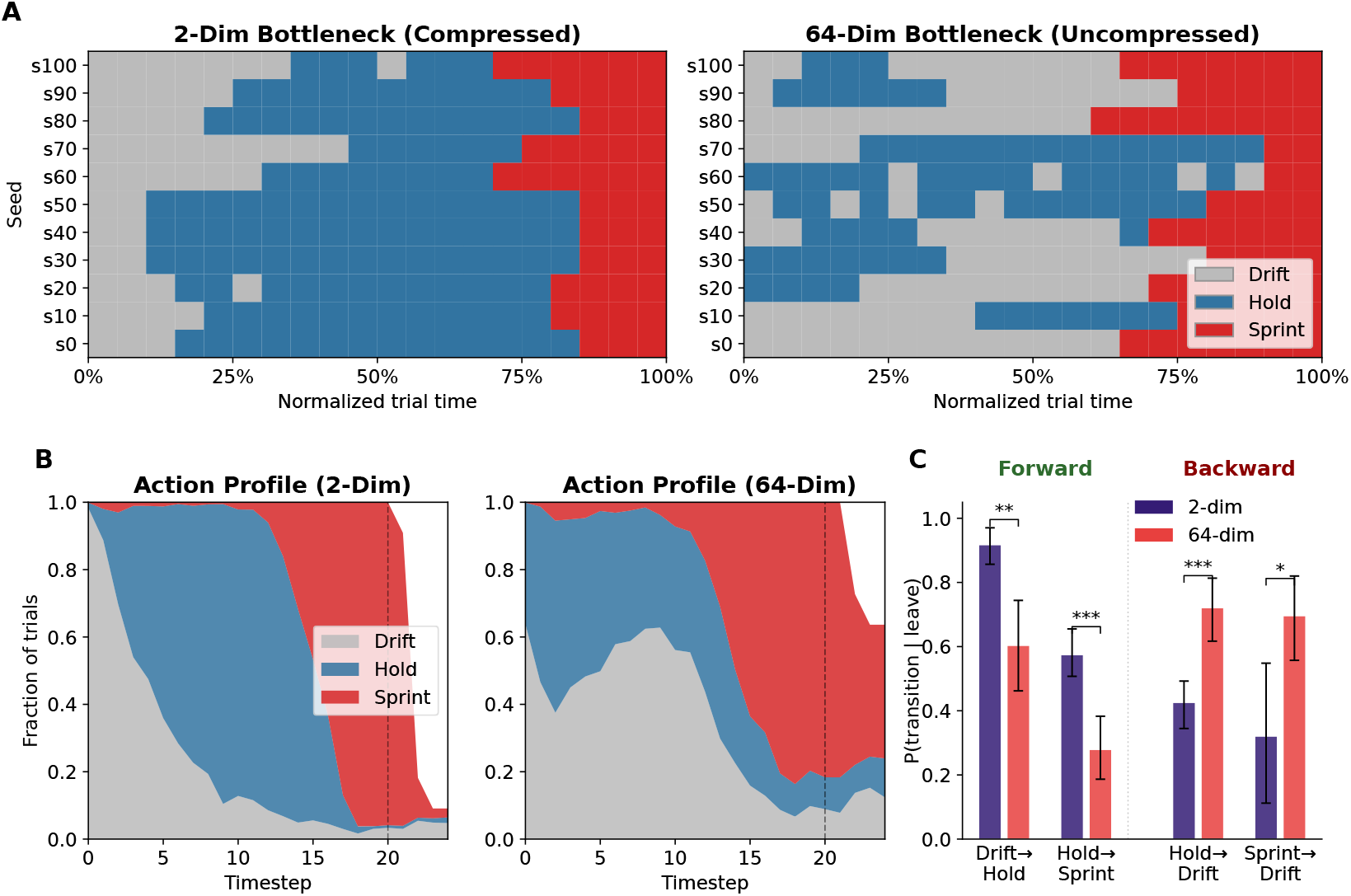
Corticostriatal compression reliably induces the Front-Back-Front motor program. **(A)** Modal action per normalized timestep for each seed, on successful trials (reward *>* 0) after time-warping to a common duration. **(B)** Mean action profile across all 11 seeds and trials. The compressed model (left) shows sharp phase boundaries; the uncompressed model (right) shows gradual, overlapping transitions. Dashed line: goal time (*t* = 20). **(C)** Conditional transition probabilities given an action switch. FBF-consistent forward transitions (Drift *→* Hold, Hold *→* Sprint) are higher in the compressed model; backward transitions (Hold *→* Drift, Sprint *→* Drift) are higher in the uncompressed model (Welch’s *t*-test; *n* = 11 seeds per condition).

**Fig. S11.**
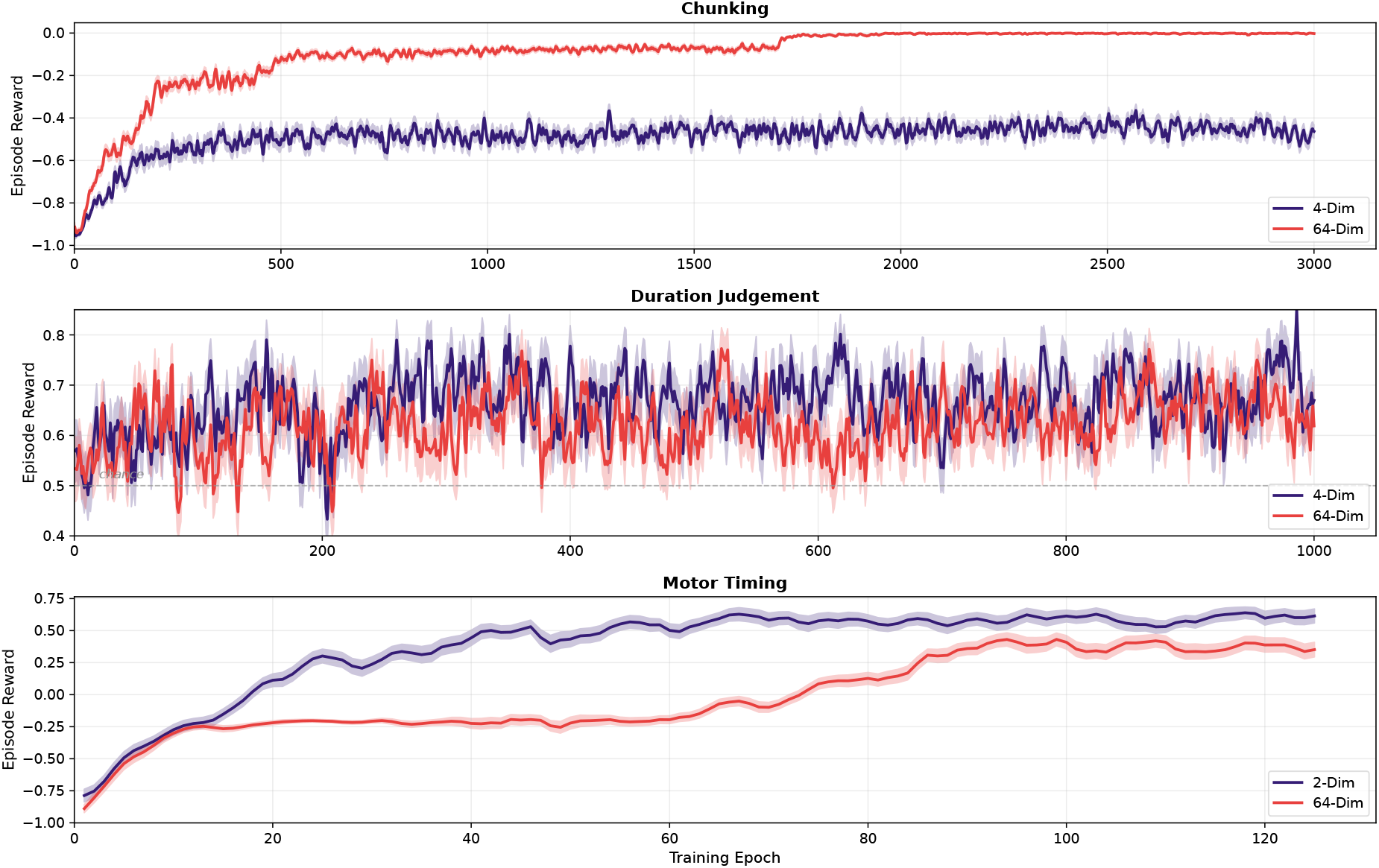
Training curves. Smoothed training curves for a representative seed (closest to the across-seed median trajectory). Shaded bands show *±*1 SEM across episodes.

### Analytical Framework: Why Compression Separates Control from Dynamics

The preceding results establish empirically that corticostriatal compression produces lower trajectory variance (Figures 3D, 5D, 7H), stronger time encoding (Figures 3D, 5C, 7H), and stereotyped behavioral programs across three tasks. We now develop an analytical framework that makes the “control versus dynamics” structure of these findings precise. The framework has two parts. First, we show that compression creates a *temporal scaffold*: by reducing the dimensionality of input-driven variability, it produces smooth, reproducible trajectories from which elapsed time is easily decoded even before any learning takes place. Second, we show how trained networks use the limited input channels that remain: by concentrating the corticostriatal projection on the slow modes of the recurrent dynamics, they ensure that each input has lasting downstream impact, maintaining *control* over extended temporal programs despite the narrow channel.

#### Linear surrogate dynamics

Consider a linear dynamical system capturing the essential structure of our corticostriatal model:

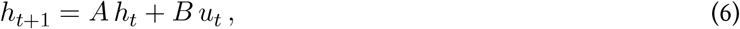

where *h*_*t*_ *∈* ℝ^*n*^ is the striatal hidden state, *A ∈* ℝ^*n×n*^ is the recurrent dynamics matrix, *B ∈* ℝ^*n×r*^ is the corticostriatal input projection with *r ≤ n*, and *u*_*t*_ ~ 𝒩 (0, *σ*^2^*I*_*r*_) is i.i.d. input noise representing the stochastic component of the cortical signal transmitted through the bottleneck. We assume *A* is *stable* (*ρ*(*A*) *<* 1, i.e. all eigenvalues lie inside the unit circle), so that the system has a well-defined steady state.

### Part I: Dynamics—Compression Creates a Temporal Scaffold

#### Steady-state covariance

Starting from *h*_0_ = 0 and iterating (6) gives

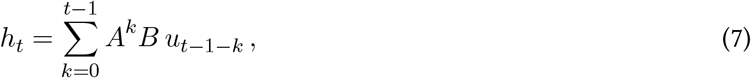

a weighted sum of past inputs: input *u*_*t*−1−*k*_ enters through *B* and is propagated *k* steps by repeated multiplication with *A*. Since each *u*_*t*_ has zero mean, so does *h*_*t*_, and its covariance is 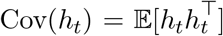. Expanding 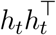 using (7) gives a double sum over all pairs of past inputs. Because the inputs are independent across time 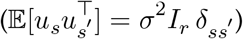, every cross-term vanishes and only the diagonal terms survive:

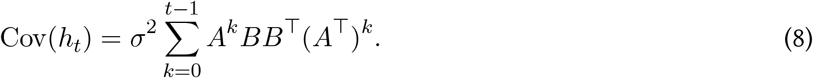

Because *A* is stable, the terms decay geometrically and the sum converges as *t → ∞* to the steady-state covariance

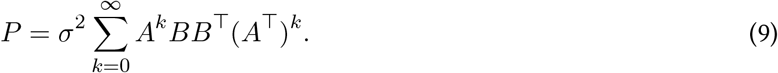

#### Trajectory variance

The diagonal entry *P*_*ii*_ is the variance of the *i*-th hidden-state component across trials: 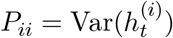. The trace sums these across all *n* dimensions, and dividing by *n* gives the mean per-dimension variance—our trajectory variance metric:

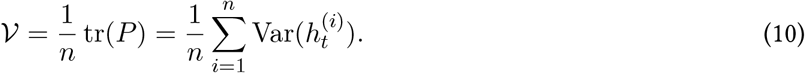

**Fig. S12.**
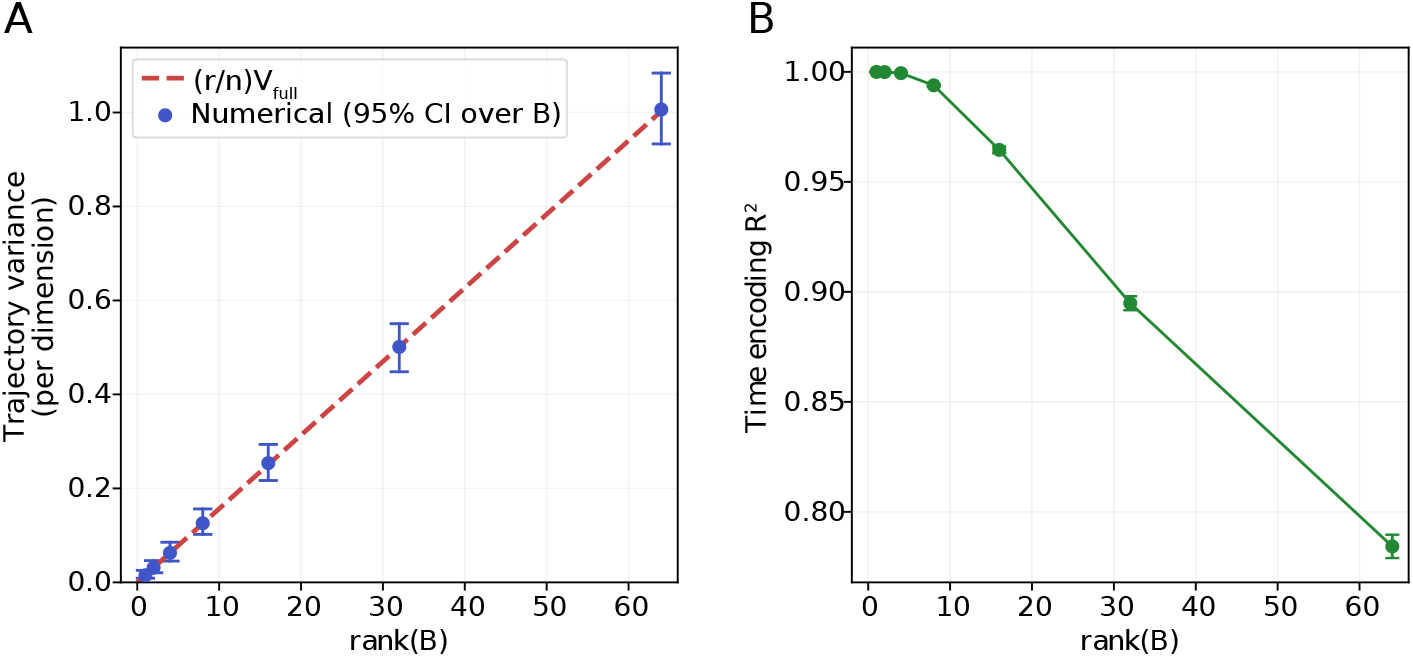
Compression creates a temporal scaffold. **(A)** Trajectory variance of a linear system *h*_*t*+1_ = *Ah*_*t*_ + *Bu*_*t*_ as a function of rank(*B*), for *n* = 64, *σ* = 0.4, *ρ*(*A*) = 0.95. Blue dots: numerical Lyapunov solution averaged over 10 random stable *A* and 20 random isotropic *B* per rank. Error bars: 95% CI over draws of *B* (*A*-variability removed by normalizing each draw by 𝒱_full_ for the same *A*). Red dashed line: analytical prediction (*r/n*) 𝒱_full_ (Proposition 0.1). The prediction, though stated in expectation, holds tightly for individual draws of *B*. **(B)** Time-encoding *R*^2^ (cross-validated Ridge regression of elapsed timestep from the hidden state) improves monotonically as rank(*B*) decreases. All conditions share the same mean trajectory; only input-noise variance differs, confirming that compression improves temporal decoding by reducing trial-to-trial variability (*n* = 300 trials, *T* = 25 timesteps, 10 random *A ×* 5 random *B* draws per rank; error bars: SEM over *B*).

#### Factoring trajectory variance through a single matrix *Q*

Substituting (9) into (10) and using the cyclic property of the trace (tr(*A*^*k*^*B · B*^⊤^(*A*^⊤^)^*k*^) = tr(*B*^⊤^(*A*^⊤^)^*k*^*A*^*k*^*B*)) to rearrange each summand:

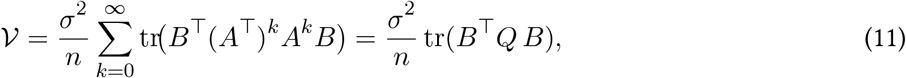

where in the last step we collected all the *A*-dependent pieces into one matrix (exchanging trace and sum is valid because every term is non-negative):

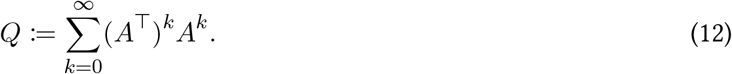

*Q* is symmetric positive semidefinite, depends only on *A*, and is the unique solution to the discrete Lyapunov equation *Q* = *I*_*n*_ + *A*^⊤^*Q A. Q* admits a direct interpretation: for any state *x*, the total energy accumulated under the autonomous dynamics is _*t≥*0_ ∥*A*^*t*^*x*∥^2^ = *x*^⊤^*Q x*. The leading eigenvectors of *Q* are therefore the directions along which state activity is most persistent.

#### Compression reduces trajectory variance: the isotropic case

Equation (11) makes the role of *B* explicit: trajectory variance is a bilinear form in *B*, weighted by *Q*. We now ask what happens for the simplest possible *B*—one whose columns are isotropically distributed in ℝ^*n*^, with no preferred direction. This is the regime of any standard weight initialization (e.g., Xavier/Glorot).

##### Proposition 0.1

(Trajectory variance scales linearly with input rank)

*Let A ∈* ℝ^*n×n*^ *be stable, and let B ∈* ℝ^*n×r*^ *be a random matrix whose columns b*_1_, …, *b*_*r*_ *each satisfy* 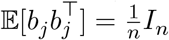 *(no assumption on the joint distribution or on Gaussianity). Then*

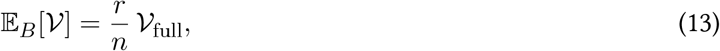

*where* 𝒱_full_ *is the trajectory variance when B* = *I*_*n*_.

*Proof*. From (11), 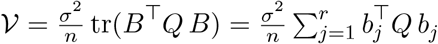, where the second equality expands the trace in terms of the columns of *B*. Taking expectations and using linearity:

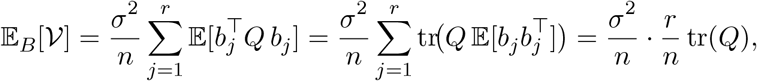

where we used the identity 𝔼 [*b*^⊤^*Mb*] = tr(*M* 𝔼 [*bb*^⊤^]) and the assumption 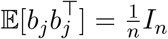. When *B* = *I*_*n*_, the same formula gives 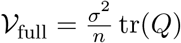, so 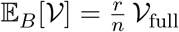.

The result is stated in expectation over *B*, but numerically the prediction holds tightly for individual draws of *B*: solving the Lyapunov equation for *P* across many random *A* and *B* matrices, the per-draw trajectory variance concentrates closely around the (*r/n*) prediction (Figure S12A, error bars show 95% CI over draws of *B*, with *A*-variability marginalized out by normalizing each draw by *V*_full_ for the same *A*).

#### The temporal scaffold

Proposition 0.1 has an immediate consequence for time encoding. The mean trajectory 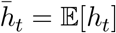 is determined by the deterministic component of cortical input and is shared across trials regardless of *r*. Trial-to-trial deviations from this mean—the noise that obscures temporal information for a linear decoder— have variance 𝒱 *≈* (*r/n*) 𝒱_full_. The signal-to-noise ratio for decoding elapsed time therefore scales as *n/r*: halving the bottleneck dimension halves the noise while leaving the signal intact (Figure S12B).

This explains a striking empirical observation: even *untrained* models with a bottleneck exhibit strong temporal structure (Figure S4). At initialization, *B* is isotropic by construction, so the proposition applies directly. Compression creates a temporal scaffold—smooth, reproducible dynamics from which elapsed time is easily decoded—as a purely geometric consequence of low input rank, before any learning takes place.

### Part II: Control—Compression Targets the Slow Modes

The temporal scaffold explains why compression reduces trajectory variance and improves time encoding. But the network must still *do* something with its limited input channels—it must exert control over the recurrent dynamics to produce task-appropriate behavior. The spectral decomposition of trajectory variance reveals how.

#### Spectral decomposition of trajectory variance

Since *Q* is symmetric, the spectral theorem gives an eigendecomposition *Q* = *U* diag(*q*_1_, …, *q*_*n*_) *U*^⊤^ with *q*_1_ *≥ q*_2_ *≥ · · · ≥ q*_*n*_ *≥* 0. Substituting into (11) and expanding (using tr(*M* ^⊤^*D M*) = Σ_*i*_ *D*_*ii*_ ∥*M*_row *i*_∥^2^ for diagonal *D*):

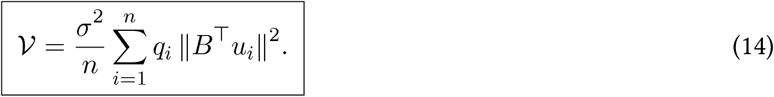

Each term has two factors:

- *q*_*i*_: the *importance weight* of direction *u*_*i*_, depending on *A*. For the specific case of normal *A, q*_*i*_ = 1*/*(1 − |*λ*_*i*_|^2^): slow modes (|*λ*_*i*_| *→* 1) have large *q*_*i*_, fast modes have *q*_*i*_ *≈* 1.
- 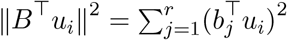: the *alignment* of the input projection with direction *u*_*i*_.

The trajectory variance is a *q*-weighted overlap of the input subspace with the eigenmodes of the recurrent dynamics. The *q*-spectra of our trained recurrent matrices are heavy-tailed: a handful of slow modes carry the bulk of the total weight (Figure S13A). This means that where *B* places its projection mass—on slow modes or fast modes—has a large effect on the resulting variance.

#### Slow modes carry the most downstream influence

The decomposition (14) has a direct dynamical reading. The total state-space energy deposited by an input pulse *u* injected through *B*, summed over all future timesteps, is

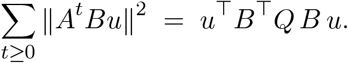

**Fig. S13.**
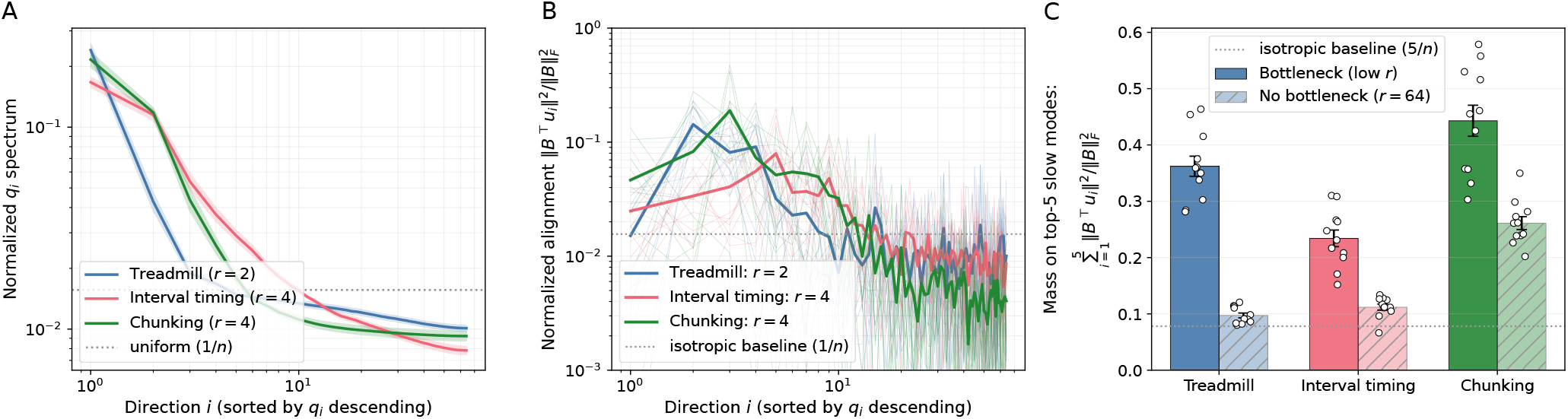
Trained corticostriatal projections target the slow modes of the recurrent dynamics. **(A)** Normalized eigenvalue spectrum of *Q* = Σ (*A*^⊤^)^*k*^*A*^*k*^ for trained recurrent weights *A* = *W*_*hg*_ (rescaled to *ρ* = 0.9) of bottleneck networks, averaged across seeds. A handful of slow modes carry the bulk of the total *q*-weight; the dotted line marks the isotropic baseline 1*/n*. **(B)** Normalized alignment 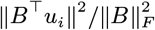 of the trained input projection *B* = *W*_*ig*_ with the eigendirections of *Q*, sorted by *q*_i_ descending, for the most-compressed condition of each task. Thin lines: individual seeds; bold: across-seed mean. The projection mass concentrates on the top several slow modes, well above the isotropic baseline. **(C)** Fraction of 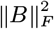 on the top-5 slow modes of *Q*, compared between bottleneck (low *r*, solid bars) and no-bottleneck (*r* = 64, hatched bars) networks. Dots: individual seeds; error bars: SEM. Dotted line: isotropic baseline 5*/n*. Bottleneck networks place systematically more of their input projection on the dominant slow modes across all three tasks, confirming that compression drives the corticostriatal projection toward the directions of striatal state space along which inputs have the most persistent downstream effect (the leading eigenvectors of Q).

Input directions aligned with the top eigenvectors of *Q* deposit disproportionately more energy per unit input, because those directions support persistent state activity over many timesteps. A *B* whose columns concentrate on the leading eigenvectors of *Q* therefore maximizes the downstream influence of each input channel: each cortical signal reverberates through the recurrent dynamics rather than being washed out.

#### Trained networks concentrate on slow modes

We tested whether trained networks exploit this structure by examining the input-to-cell-update weight matrices *B* = *W*_*ig*_ and recurrent-to-cell-update matrices *A* = *W*_*hg*_ of our LSTMs.^1^. The alignment profile of trained bottleneck networks confirmed the prediction: ∥*B*^⊤^*u*_*i*_ ∥^2^ was elevated 3–12*×* above the isotropic baseline 1*/n* for the top several slow modes and dropped below base*∥*line in the fast directions (Figure S13B). The effect was compression-specific: aggregating the mass of *B* on the top five slow modes of *Q*, bottleneck networks consistently placed 3–4*×* more mass there than their matched full-rank counterparts on every task (Figure S13C).

This slow-mode concentration is the trained network’s solution to the control problem posed by the bottleneck. With only *r* ≪ *n* input channels available, the corticostriatal projection concentrates on the directions where input has the most persistent downstream effect (slowly decaying eigenmodes). Each cortical signal is thereby amplified across multiple timesteps, enabling control over extended temporal programs despite the narrow channel. This is the “control” complement to the “dynamics” story of Part I: compression creates a stable temporal scaffold by reducing input-driven variability, and the network maintains task competence by routing its limited input through long-range modes.

## Acknowledgments

S.K is currently supported by a Leon Levy Fellowship by the New York Academy of Sciences. J.H is supported by VA CSR&D award 1I01 CX002872.

## Footnotes

1 Our analysis uses the recurrent-to-cell-update weight matrix *W* _*hg*_, rescaled to spectral radius *ρ* = 0.9, as a linear surrogate for the full LSTM dynamics. To validate this choice, we also computed the effective Jacobian using the mean-field gate-modulation framework of Can et al. (2020), which incorporates all four gate weight matrices and biases without requiring task inputs. The resulting alignment profiles were nearly identical (e.g., top-5 slow-mode fraction of 0.38 vs. 0.36 for treadmill bottleneck networks), confirming that gating modulates the spectral scale of *W*_*hg*_ without reorganizing its eigenstructure. We report the simpler *W*_*hg*_-based analysis. conclusions are robust to the choice of *ρ ∈* [0.8, 0.95].

## References

Alexander, G. E., DeLong, M. R., & Strick, P. L. (1986). Parallel organization of functionally segregated circuits linking basal ganglia and cortex. Annual review of neuroscience, 9(1), 357–381.

Arabzadeh, E., Panzeri, S., & Diamond, M. E. (2004). Whisker vibration information carried by rat barrel cortex neurons. Journal of Neuroscience, 24(26), 6011–6020.

Bar-Gad, I., Havazelet-Heimer, G., Goldberg, J., Ruppin, E., & Bergman, H. (2003). Reinforcement driven dimensionality reduction as a model for information processing in the basal ganglia. Hebrew University of Jerusalem.

Bar-Gad, I., Morris, G., & Bergman, H. (2003). Information processing, dimensionality reduction and reinforcement learning in the basal ganglia. Progress in neurobiology, 71(6), 439–473.

Botvinick, M., & Plaut, D. C. (2004). Doing without schema hierarchies: a recurrent connectionist approach to normal and impaired routine sequential action. Psychological review, 111(2), 395.

Buonomano, D. V., & Maass, W. (2009). State-dependent computations: spatiotemporal processing in cortical networks. Nature Reviews Neuroscience, 10(2), 113–125.

Can, T., Krishnamurthy, K., & Schwab, D. J. (2020). Gating creates slow modes and controls phase-space complexity in grus and lstms. In Mathematical and scientific machine learning (pp. 476–511).

Dhawale, A. K., Wolff, S. B., Ko, R., & Ölveczky, B. P. (2021). The basal ganglia control the detailed kinematics of learned motor skills. Nature neuroscience, 24(9), 1256–1269.

Dominey, P. F. (1995). Complex sensory-motor sequence learning based on recurrent state representation and reinforcement learning. Biological cybernetics, 73(3), 265–274.

Doya, K. (2000). Complementary roles of basal ganglia and cerebellum in learning and motor control. Current opinion in neurobiology, 10(6), 732–739.

Eichenbaum, H. (2014). Time cells in the hippocampus: a new dimension for mapping memories. Nature Reviews Neuroscience, 15(11), 732–744.

Gouvêa, T. S., Monteiro, T., Motiwala, A., Soares, S., Machens, C., & Paton, J. J. (2015). Striatal dynamics explain duration judgments. Elife, 4, e11386.

Graybiel, A. M. (1998). The basal ganglia and chunking of action repertoires. Neurobiology of learning and memory, 70(1-2), 119–136.

Haber, S. N. (2003). The primate basal ganglia: parallel and integrative networks. Journal of chemical neuroanatomy, 26(4), 317–330.

Haber, S. N., Fudge, J. L., & McFarland, N. R. (2000). Striatonigrostriatal pathways in primates form an ascending spiral from the shell to the dorsolateral striatum. Journal of Neuroscience, 20(6), 2369–2382.

Haimerl, C., Rodrigues, F. S., & Paton, J. J. (2025). Time, control, and the nervous system. Annual Review of Neuroscience, 48.

Hidalgo-Balbuena, A. E., Luma, A. Y., Pimentel-Farfan, A. K., Peña-Rangel, T., & Rueda-Orozco, P. E. (2019). Sensory representations in the striatum provide a temporal reference for learning and executing motor habits. Nature communications, 10(1), 4074.

Hochreiter, S., & Schmidhuber, J. (1997). Long short-term memory. Neural computation, 9(8), 1735–1780.

Humphries, M. D. (2025). The computational bottleneck of basal ganglia output (and what to do about it). eneuro, 12(4).

Jeffery, K. J. (2007). Integration of the sensory inputs to place cells: what, where, why, and how? Hippocampus, 17 (9), 775–785.

Jin, D. Z., Fujii, N., & Graybiel, A. M. (2009). Neural representation of time in cortico-basal ganglia circuits. Proceedings of the National Academy of Sciences, 106(45), 19156–19161.

Jin, X., & Costa, R. M. (2015). Shaping action sequences in basal ganglia circuits. Current opinion in neurobiology, 33, 188–196.

Kincaid, A. E., Zheng, T., & Wilson, C. J. (1998). Connectivity and convergence of single corticostriatal axons. Journal of Neuroscience, 18(12), 4722–4731.

Kingma, D. P., & Ba, J. (2014). Adam: A method for stochastic optimization. arXiv preprint arXiv:1412.6980.

Kita, H., & Kitai, S. (1988). Glutamate decarboxylase immunoreactive neurons in rat neostriatum: their morphological types and populations. Brain research, 447 (2), 346–352.

Klaus, A., Martins, G. J., Paixao, V. B., Zhou, P., Paninski, L., & Costa, R. M. (2017). The spatiotemporal organization of the striatum encodes action space. Neuron, 95(5), 1171–1180.

Lai, L., & Gershman, S. J. (2021). Policy compression: An information bottleneck in action selection. In Psychology of learning and motivation (Vol. 74, pp. 195–232). Elsevier.

Lai, L., Huang, A. Z., & Gershman, S. J. (2025). Action chunking as conditional policy compression. Cognition, 264, 106201.

Lanciego, J. L., Luquin, N., & Obeso, J. A. (2012). Functional neuroanatomy of the basal ganglia. Cold Spring Harbor perspectives in medicine, 2(12), a009621.

Lee, K., Bakhurin, K. I., Claar, L. D., Holley, S. M., Chong, N. C., Cepeda, C., … Masmanidis, S. C. (2019). Gain modulation by corticostriatal and thalamostriatal input signals during reward-conditioned behavior. Cell reports, 29(8), 2438–2449.

Li, M., Jensen, K. T., Zhang, Q., Lu, Q., & Mattar, M. G. (2025). A neural network model of free recall learns multiple memory strategies. bioRxiv, 2025–09.

Liang, E., Liaw, R., Nishihara, R., Moritz, P., Fox, R., Goldberg, K., … Stoica, I. (2018). Rllib: Abstractions for distributed reinforcement learning. In International conference on machine learning (pp. 3053–3062).

Logiaco, L., Abbott, L., & Escola, S. (2021). Thalamic control of cortical dynamics in a model of flexible motor sequencing. Cell reports, 35(9).

Mandelbaum, G., Taranda, J., Haynes, T. M., Hochbaum, D. R., Huang, K. W., Hyun, M., … others (2019). Distinct cortical-thalamic-striatal circuits through the parafascicular nucleus. Neuron, 102(3), 636–652.

Markowitz, J. E., Gillis, W. F., Jay, M., Wood, J., Harris, R. W., Cieszkowski, R., … others (2023). Spontaneous behaviour is structured by reinforcement without explicit reward. Nature, 614(7946), 108–117.

Martiros, N., Burgess, A. A., & Graybiel, A. M. (2018). Inversely active striatal projection neurons and interneurons selectively delimit useful behavioral sequences. Current Biology, 28(4), 560–573.

McElvain, L. E., Chen, Y., Moore, J. D., Brigidi, G. S., Bloodgood, B. L., Lim, B. K., … Kleinfeld, D. (2021). Specific populations of basal ganglia output neurons target distinct brain stem areas while collateralizing throughout the diencephalon. Neuron, 109(10), 1721–1738.

Medina, J. F., Garcia, K. S., Nores, W. L., Taylor, N. M., & Mauk, M. D. (2000). Timing mechanisms in the cerebellum: testing predictions of a large-scale computer simulation. Journal of Neuroscience, 20(14), 5516–5525.

Mizes, K. G., Lindsey, J., Escola, G. S., & Ölveczky, B. P. (2023). Dissociating the contributions of sensorimotor striatum to automatic and visually guided motor sequences. Nature Neuroscience, 26(10), 1791–1804.

Mizes, K. G., Lindsey, J., Escola, G. S., & Ölveczky, B. P. (2024). The role of motor cortex in motor sequence execution depends on demands for flexibility. Nature neuroscience, 27 (12), 2466–2475.

Monteiro, T., Rodrigues, F. S., Pexirra, M., Cruz, B. F., Gonçalves, A. I., Rueda-Orozco, P. E., & Paton, J. J. (2023). Using temperature to analyze the neural basis of a time-based decision. Nature neuroscience, 26(8), 1407–1416.

Mountcastle, V. B., Talbot, W. H., Sakata, H., & Hyvärinen, J. (1969). Cortical neuronal mechanisms in flutter-vibration studied in unanesthetized monkeys. neuronal periodicity and frequency discrimination. Journal of neurophysiology, 32(3), 452–484.

Murray, J. M., & Escola, G. S. (2017). Learning multiple variable-speed sequences in striatum via cortical tutoring. Elife, 6, e26084.

Nassar, M. R., Helmers, J. C., & Frank, M. J. (2018). Chunking as a rational strategy for lossy data compression in visual working memory. Psychological review, 125(4), 486.

Norman, D. A. (1981). Categorization of action slips. Psychological review, 88(1), 1.

Pasupathy, A., & Miller, E. K. (2005). Different time courses of learning-related activity in the prefrontal cortex and striatum. Nature, 433(7028), 873–876.

Rodrigues, F. S., Monteiro, T., Motiwala, A., & Paton, J. J. (2024). The dorsolateral striatum encodes a temporal basis for the organization of behavior. Neuron, 112(22), 3675–3677.

Rueda-Orozco, P. E., & Robbe, D. (2015). The striatum multiplexes contextual and kinematic information to constrain motor habits execution. Nature neuroscience, 18(3), 453–460.

Schulman, J., Wolski, F., Dhariwal, P., Radford, A., & Klimov, O. (2017). Proximal policy optimization algorithms. arXiv preprint arXiv:1707.06347.

Soni, A., & Frank, M. J. (2025). Adaptive chunking improves effective working memory capacity in a prefrontal cortex and basal ganglia circuit. Elife, 13, RP97894.

Thorn, C. A., Atallah, H., Howe, M., & Graybiel, A. M. (2010). Differential dynamics of activity changes in dorsolateral and dorsomedial striatal loops during learning. Neuron, 66(5), 781–795.

Toso, A., Reinartz, S., Pulecchi, F., & Diamond, M. E. (2021). Time coding in rat dorsolateral striatum. Neuron, 109(22), 3663–3673.

Tsao, A., Yousefzadeh, S. A., Meck, W. H., Moser, M.-B., & Moser, E. I. (2022). The neural bases for timing of durations. Nature reviews neuroscience, 23(11), 646–665.

Wagner, M. J., Kim, T. H., Kadmon, J., Nguyen, N. D., Ganguli, S., Schnitzer, M. J., & Luo, L. (2019). Shared cortex-cerebellum dynamics in the execution and learning of a motor task. Cell, 177 (3), 669–682.

Wang, J. X., Kurth-Nelson, Z., Kumaran, D., Tirumala, D., Soyer, H., Leibo, J. Z., … Botvinick, M. (2018). Prefrontal cortex as a meta-reinforcement learning system. Nature neuroscience, 21(6), 860–868.

Wilson, C. J., & Groves, P. M. (1980). Fine structure and synaptic connections of the common spiny neuron of the rat neostriatum: a study employing intracellular injection of horseradish peroxidase. Journal of Comparative Neurology, 194(3), 599–615.

Yin, H. H., Knowlton, B. J., & Balleine, B. W. (2004). Lesions of dorsolateral striatum preserve outcome expectancy but disrupt habit formation in instrumental learning. European journal of neuroscience, 19(1), 181–189.

